# Lamellipodin tunes cell migration by stabilizing protrusions and promoting adhesion formation

**DOI:** 10.1101/777326

**Authors:** Georgi Dimchev, Behnam Amiri, Ashley C. Humphries, Matthias Schaks, Vanessa Dimchev, Theresia E.B. Stradal, Jan Faix, Matthias Krause, Michael Way, Martin Falcke, Klemens Rottner

## Abstract

Efficient migration on adhesive surfaces involves the protrusion of lamellipodial actin networks and their subsequent stabilization by nascent adhesions. The actin binding protein lamellipodin (Lpd) is thought to play a critical role in lamellipodium protrusion, by delivering Ena/VASP proteins onto the growing plus ends of actin filaments and by interacting with the WAVE regulatory complex (WRC), an activator of the Arp2/3 complex, at the leading edge. Using B16-F1 melanoma cell lines, we demonstrate that genetic ablation of Lpd compromises protrusion efficiency and coincident cell migration without altering essential parameters of lamellipodia, including their maximal rate of forward advancement and actin polymerization. We also confirmed lamellipodia and migration phenotypes with CRISPR/Cas9-mediated Lpd knockout Rat2 fibroblasts, excluding cell type-specific effects. Moreover, computer-aided analysis of cell edge morphodynamics on B16-F1 cell lamellipodia revealed that loss of Lpd correlates with reduced temporal protrusion maintenance as a prerequisite of nascent adhesion formation. We conclude that Lpd optimizes protrusion and nascent adhesion formation by counteracting frequent, chaotic retraction and membrane ruffling.

**Summary statement:** We describe how genetic ablation of the prominent actin- and VASP-binding protein lamellipodin combined with software-aided protrusion analysis uncovers mechanistic insights into its cellular function during cell migration.

## INTRODUCTION

The protrusion of lamellipodia and filopodia at the plasma membrane is crucial for various processes ranging from migration of individual, mesenchymal or tumour cells to neuronal growth cone advance or epithelial zippering during adherens junction formation (Bachir et al., 2017; Omotade et al., 2017; Rottner et al., 2017). Lamellipodin (Lpd), a member of the Mig10/RIAM/Lpd family of adaptor proteins (MRL), localizes at the edges or tips of protruding lamellipodia and filopodia and interacts with multiple factors involved in the regulation of assembly (Colo et al., 2012; Krause and Gautreau, 2014). Lpd supports lamellipodia protrusion and cell migration, to promote cancer cell invasion and neural crest migration. It also supports neuronal morphogenesis, endocytosis and localises to the interface between vaccinia virus and their actin tails (Bodo et al., 2017; Boucrot et al., 2015; Carmona et al., 2016; Chan Wah Hak et al., 2018; Hansen and Mullins, 2015; Krause et al., 2004; Law et al., 2013; Vehlow et al., 2013). All these functions are mediated by individual Lpd domains (Fig. S1), but we still need a better understanding of how Lpd functions via its respective binding partners. For example, plasma membrane localisation of Lpd is clearly facilitated by its pleckstrin homology (PH) domain, which interacts with PI(3,4)P2 membrane phospholipids (Bae et al., 2014; Krause et al., 2004; Smith et al., 2010a). Nevertheless, actin filament binding through C-terminal sequences lacking the PH-domain is necessary and sufficient for Lpd enrichment at the lamellipodium edge (Hansen and Mullins, 2015).

Lpd can regulate lamellipodia formation and cell migration through interactions with WAVE Regulatory Complex (WRC) (Law et al., 2013), a potent activator of the Arp2/3 complex (Rottner and Schaks, 2019; Schaks et al., 2018), or with both WRC and Ena/VASP family proteins, depending on the cell system and conditions (Carmona et al., 2016; Krause and Gautreau, 2014; Krause et al., 2004). Lpd binds directly to the SH3-domain of the WRC subunit Abi, and this interaction is positively regulated by Rac binding to the Ras association (RA) domain of Lpd as well as by c-Abl and c-Src-dependent phosphorylation (Law et al., 2013). In contrast, the association between Lpd and Ena/VASP proteins is specifically regulated by Abl but not Src family kinases and occurs through its C-terminal Ena/VASP homology 1- (EVH1-) binding sites (Hansen and Mullins, 2015; Krause et al., 2004; Michael et al., 2010). The potential relevance of this interaction to lamellipodial dynamics is supported by the prominent accumulation of both Lpd and Ena/VASP proteins at the edges of protruding lamellipodia (Hansen and Mullins, 2015; Krause et al., 2004; Rottner et al., 1999a). Mechanistically, the actin filament binding activity of Lpd, which can occur independently of Ena/VASP binding, was proposed to tether the latter to growing plus ends, thereby increasing their processive polymerase activity (Hansen and Mullins, 2015).

Lpd promotes migration processes in diverse developmental processes, ranging from neural crest-derived melanoblast migration in mice to border cell migration in *Drosophila* (Law et al., 2013). Through its Ena/VASP and WRC interactions, Lpd also increases cancer cell invasion, and its increased expression in breast cancer samples correlates with poor patient prognosis (Carmona et al., 2016). At a cellular level, Lpd overexpression increases the speed of lamellipodia protrusion, while its knockdown by RNA interference (RNAi) or conditional genetic knockout impairs their formation (Carmona et al., 2016; Krause et al., 2004; Law et al., 2013). However, the consequences of permanent loss of Lpd function by genetic ablation in growing cell lines has not been studied. Lpd has also been implicated in various additional processes either involving or at least impacting on actin dynamics. These processes include additional types of protrusion, such as those mediating cell-to-cell spreading of pathogenic *Listeria monocytogenes* (Wang et al., 2015), but also integrin activation through its binding to talin (Lagarrigue et al., 2015; Lee et al., 2009; Watanabe et al., 2008) or the fast endophilin-mediated endocytosis (FEME) pathway (Boucrot et al., 2015; Chan Wah Hak et al., 2018). How Lpd is recruited to and regulated within these distinct subcellular structures, remains to be investigated.

In this study, we generated B16-F1 mouse melanoma cell lines genetically disrupted for Lpd by CRISPR/Cas9, to precisely determine the consequences of the complete loss of Lpd function. We also describe a novel cell edge analysis workflow that employs edge detection via ImageJ with quantitative morphodynamic analysis to retrieve a large number of parameters to describe complex cellular protrusion and retraction phenotypes. Our analysis reveals that while not essential, Lpd optimizes leading edge protrusions, but without qualitatively or quantitatively affecting the rates of lamellipodial actin network polymerization. Instead, Lpd loss of function leads to changes in nascent adhesion number and distribution. We conclude that Lpd contributes to the efficiency of cell migration by helping to coordinate actin dynamics with cell-substratum adhesion.

## RESULTS

### Genetic deletion of reduces lamellipodial protrusion and rates of migration

To investigate the effects of eliminating Lpd expression on lamellipodial protrusion, we disrupted the *Lpd* gene in murine B16-F1 melanoma cells using CRISPR/Cas9. Multiple B16-F1 cell clones were isolated from the CRISPR/Cas9 transfection pool, of which 3 clones were randomly selected for further characterization (termed Lpd KO#3, #8 and #10). Sequences of disrupted alleles present in each clone including respective stop codon positions as well as location of CRISPR/Cas9-induced frameshifts relative to prominent Lpd domains are shown in Fig. S1A-C. Immunoblot analyses with three different antibodies confirmed the absence of Lpd expression in the selected cell lines (Fig. 1A, Fig. S1D, E). Individual clones harboured distinct numbers of disrupted, but no wildtype alleles (Fig. S1A, B and Fig. 1B). Stop codons in disrupted alleles were induced just upstream of the RA-domain of Lpd, so a potential truncated protein – if at all present - would be devoid of all major regulatory domains of Lpd (Fig. S1C). Immunofluorescence analysis with an antibody specific for the C-terminus of Lpd revealed that the localization of Lpd at lamellipodia edges seen in wildtype B16-F1 cells (Krause et al., 2004) was absent in the three *Lpd* gene-disrupted cell lines (Fig. S1F). We also explored the expression of RIAM/PREL-1, the second member of the MRL (Mig-10, Riam and lamellipodin) family, but consistent with previous observations (Jenzora et al., 2005), the protein was undetectable by Western Blotting in B16-F1 wildtype cells in contrast to NIH 3T3 fibroblasts (Fig. S1G). RIAM/PREL-1 was also not upregulated in our Lpd-deficient cell lines (Fig. S1H).

**Fig. 1.**
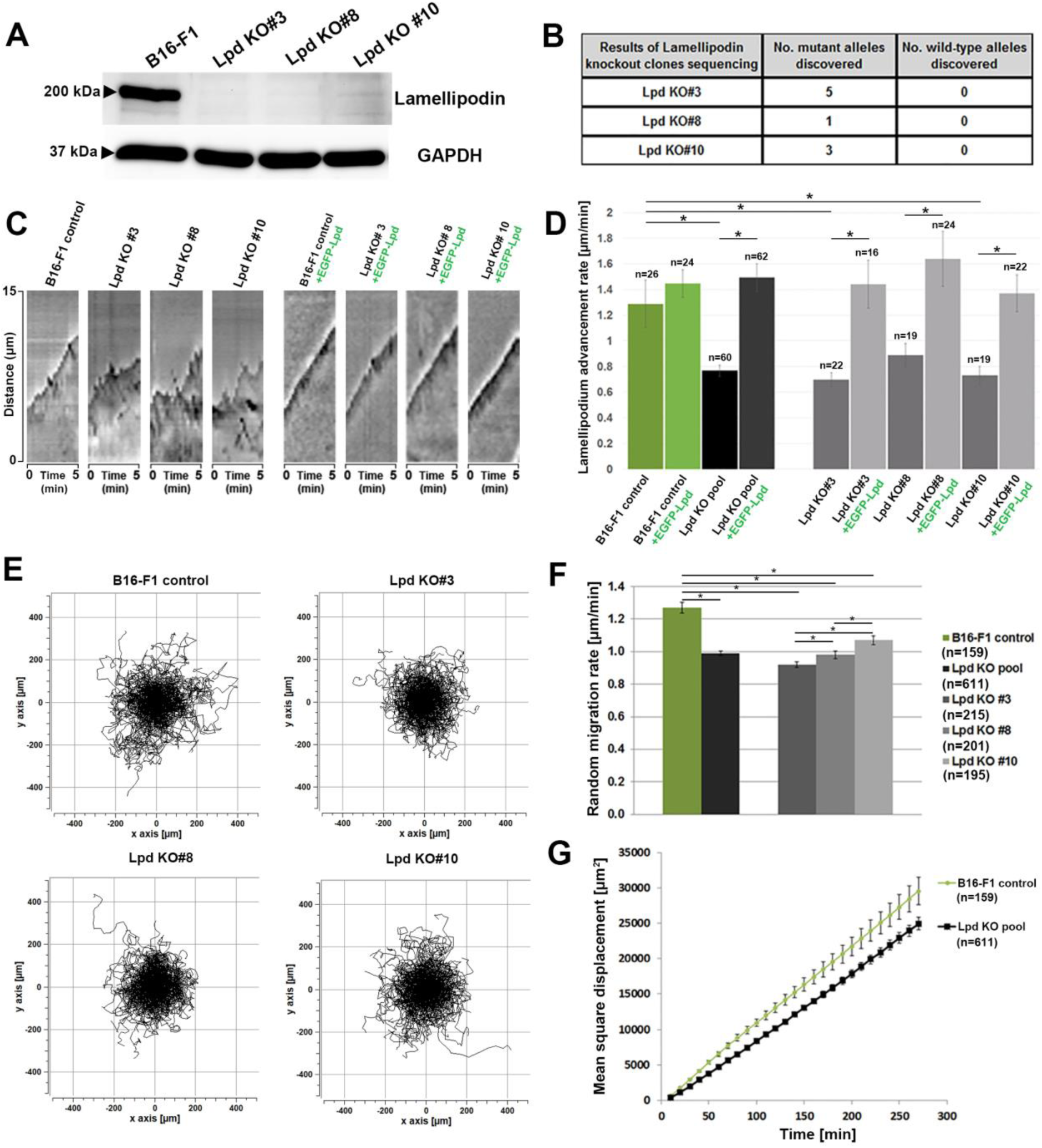
Lamellipodin (Lpd) deletion in B16-F1 cells reduces rates of lamellipodial protrusion and random migration. (A) Immunoblot analysis using an antibody recognizing the C-terminus of Lpd demonstrates the Lpd KO#3, #8 and #10 cell lines do not contain detectable levels of the protein. GAPDH represents a loading control (B) Table summarizes DNA sequencing of the genomic target in Lpd KO#3, #8 and #10 cell lines from at least 20 sequencing reactions each. (C) Kymographs of protruding lamellipodia of B16-F1 wildtype and Lpd-KO cells or either cell type expressing EGFP-Lpd, as indicated over a 5 min interval. (D) Quantitation of average lamellipodial advancement rates using kymography as raw data. (E) Trajectory plots derived from manual tracking of individually migrating cells of the indicated genotypes on laminin. (F) Quantitation of the data shown in E. (G) Mean square displacement over time of B16-F1 control and the Lpd KO pool derived from data in E. All quantified data are displayed as arithmetic means +/− SEM, with asterisks above bar charts indicating statistically significant differences between designated groups, * = p<0.05, and n = number of cells analysed.

Interestingly, we observed that lamellipodia were still able to form in the absence of Lpd in our B16-F1 CRISPR KO lines (Fig. S1F, insets, see also below). To assess the impact of the loss of Lpd on lamellipodial dynamics, we initially performed a manual, kymography-based analysis, determining the protrusion and retraction behaviour in B16-F1 wildtype and Lpd KO clones with or without expression of EGFP-Lpd over 5 minutes (Fig. 1C). This analysis revealed that protrusions were more irregular and less efficient (Fig. 1C, D). Moreover, the reduction of protrusion was fully rescued by expression of EGFP-Lpd (Fig. 1C, D), confirming the observed phenotypes are only dependent on Lpd. EGFP-Lpd expression in wildtype cells did not increase lamellipodial protrusion rates in a statistically significant fashion (Fig. 1D), suggesting that Lpd function in protrusions is saturated in this cell type. Since the efficiency of cell migration is highly dependent on lamellipodia in B16-F1 cells (Dolati et al., 2018; Schaks et al., 2018), we also examined random cell migration of our cell lines on laminin (Movie 1). Individual cell migration trajectory plots and corresponding quantifications reveals a moderate, but statistically significant reduction in random migration rates in the absence of Lpd (by ∼22%, Fig. 1E, F). This was also reflected by reduced mean square displacement of Lpd KO cells compared to wildtype controls (Fig. 1G). Together, our results reveal that B16-F1 cells lacking Lpd have reduced, but not eliminated lamellipodia protrusion activity and decreased cell migration efficiency.

To confirm these results in a different cell type and by an alternative, more transient approach, we transfected Rat2 fibroblasts with our CRISPR/Cas9-constructs that also target rat Lpd (Fig. 2A). Rat2 cells constitute the only fibroblasts in our hands lacking detectable levels of RIAM/PREL1 (Jenzora et al., 2005), excluding interference of the latter with any Lpd loss of function phenotypes. Interestingly, and although analysed as bulk population instead of single cell clones, phenotypes were highly comparable to our results obtained with individually isolated B16-F1 clones. Essentially, the vast majority of Lpd-deficient cells displayed lamellipodia that were indistinguishable in appearance to control cell lamellipodia (Fig. 2B). Moreover, the reduction of random cell migration in the transiently CRISPR/Cas9-treated population as compared to controls was also highly reminiscent of what was observed in B16-F1 Lpd KO clones (reduction by ∼14.5%, Fig. 2C, D, compare with Fig. 1E, F).

**Fig. 2.**
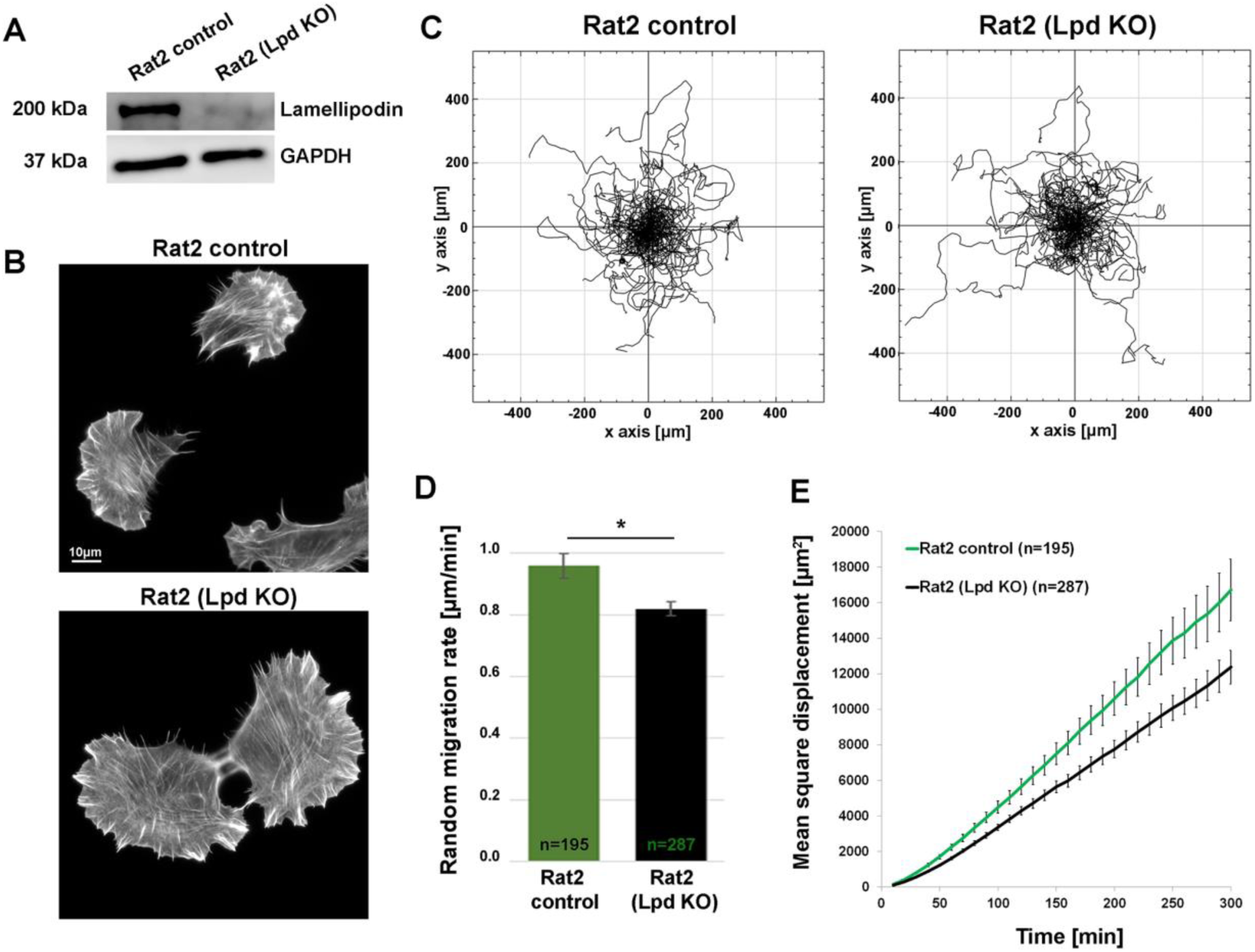
Lamellipodin (Lpd) deletion in Rat2 cells reduces rates of random migration. (A) Immunoblot analysis using an antibody recognizing the C-terminus of Lpd demonstrates that a Rat2 cell population transfected with a CRISPR/Cas9-construct targeting the Lpd gene and selected for puromycin resistance gene expression from the same plasmids (see Methods) displays hardly detectable Lpd expression (Lpd KO). Cells going through identical procedure but with non-targeting CRISPR/Cas9 vector were used as control cell population, and GAPDH was loading control. (B) Representative images of Rat2 cell populations treated as indicated (control *versus* Lpd KO) and migrating on laminin, fixed and stained for F-actin with phalloidin. (C) Trajectory plots derived from manual tracking of individually migrating cells on laminin of cell populations as indicated. (D) Quantitation of the data shown in C. (E) Mean square displacement over time of Rat2 control and Lpd KO cell populations derived from data in C. All quantified data are displayed as arithmetic means +/− SEM, with asterisk above bar charts indicating statistically significant differences between designated groups, * = p<0.05, and n = number of cells analysed.

### Loss of Lpd induces a shift towards a chaotic lamellipodial phenotype

To uncover the basis for reduced lamellipodial protrusion upon Lpd deletion, we explored lamellipodial protrusion patterns in our B16-F1 cell lines in further detail. Not surprisingly, we observed that at any given time point, lamellipodial protrusion phenotypes can differ significantly between cells within the same population. Lamellipodial dynamics also display heterogeneous patterns over time, even at the single cell level (Wang et al., 2018). To deal with the variability inherent in the system and to unambiguously display the heterogeneity in morphodynamic cell edge behaviours, we used semi-automatic cell edge tracking in cell lines expressing plasma membrane-targeted CAAX-EGFP. To achieve this, we used an ImageJ-based plug-in called JFilament (Smith et al., 2010b), followed by quantitative analysis of lamellipodial morphodynamics (see Methods). The spatiotemporal evolution of tracked lamellipodial contours over representative time periods of 2 minutes reveals there are three basic types of lamellipodial protrusion classes designated as smooth, intermediate and chaotic within B16-F1 control and Lpd knockout cell populations (Fig. 3A). Lamellipodia corresponding to the smooth phenotype protruded in a steady and persistent manner. Lamellipodia of intermediate phenotype protruded in a fluctuating manner, and chaotic lamellipodia underwent rapid cycles of protrusion and retraction, with the cell edge frequently folding backwards. Velocity and curvature maps allow clear visualization of the spatiotemporal differences of tracked cell edges between the three protrusion classes, with spatial fluctuations in both parameters clearly increasing from smooth to chaotic over time (Fig. 3B, C and Movies 2 and 3).

**Fig. 3.**
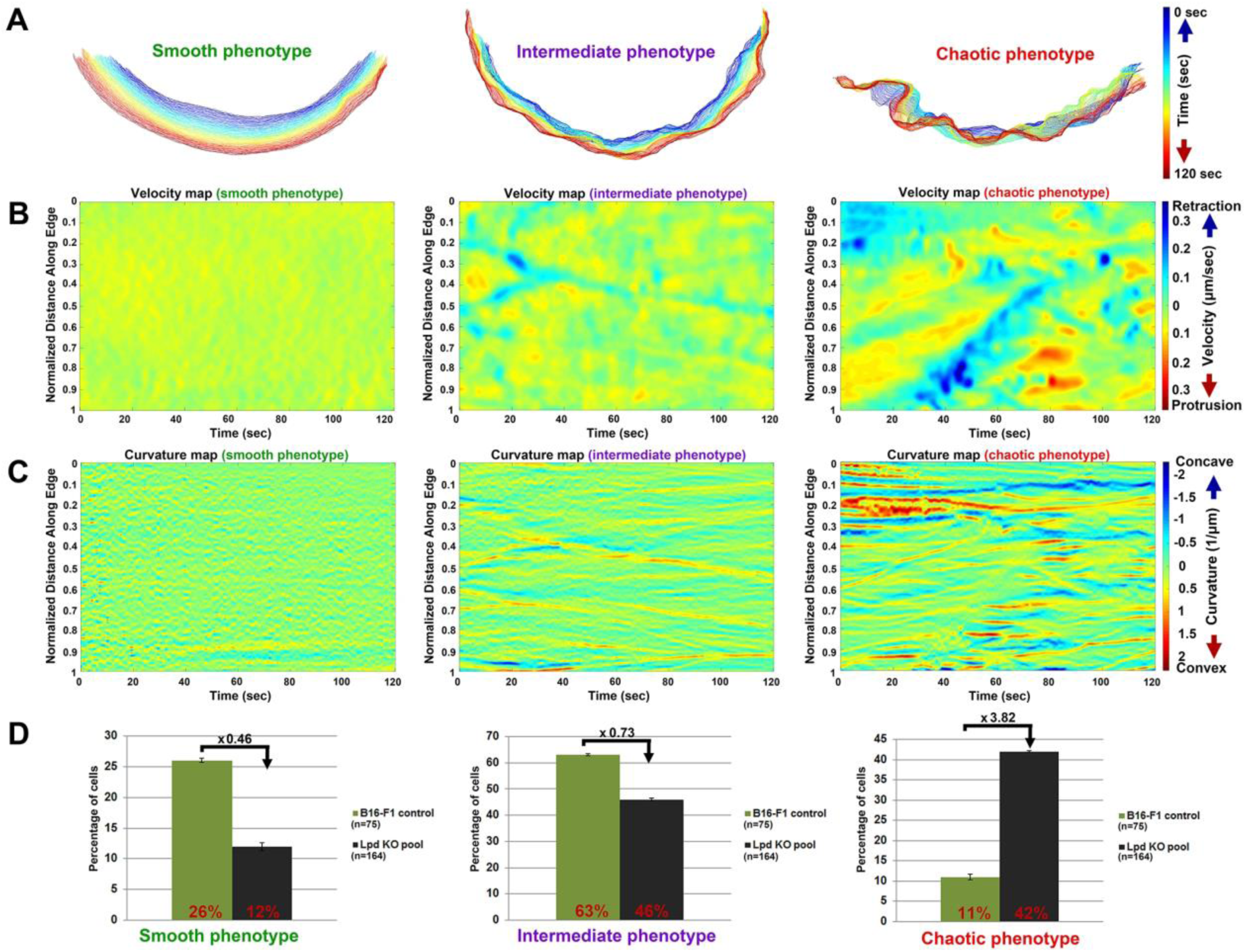
Deletion of Lpd in B16-F1 cells changes the balance between lamellipodial protrusion phenotypes. (A) Illustration of three distinct types of lamellipodial protrusion phenotypes in B16-F1 wildtype cells using computer-aided analyses of cell edge dynamics. Phenotypes are represented by the spatiotemporal evolution of measured cell edge contours, with time increasing from blue to red over 120 seconds. (B) Example velocity maps, representing smooth, intermediate and chaotic lamellipodial phenotypes, as derived by plotting the velocities of individual points into a 2D-coordinate system, with normalized distance of points along the edge on the y-axis and time on the x-axis. Retraction velocities are indicated in blue, and protrusion velocities in red (µm/sec). (C) Example curvature maps for each protrusion phenotype, as indicated, and individually derived by plotting local curvatures of individual points into a 2D-coordinate system, with normalized distance of points along the edge on the y-axis and time on the x-axis. Concave and convex curvatures are indicated in blue and red, respectively. (D) Quantification of the fraction of cells with indicated genotypes displaying each lamellipodial phenotype (smooth, intermediate or chaotic). Lpd knockout cells represent a pool of the three knockout clones.

Since we observed that Lpd knockout clones have a reduced average rate of lamellipodial protrusion, we investigated whether the fraction of cells for a given protrusion class is altered compared to controls. Analysis of movies reveals that the fraction of cells with smooth and intermediate protrusions were significantly reduced in Lpd KO cells (> two times for smooth, Fig. 3D), whereas the chaotic class increased by a factor of 3.82 (Fig. 3D, also see Movie 1). These data show there is a clear correlation between Lpd expression and lamellipodial protrusion stability.

### Lpd loss impacts on multiple morphodynamic parameters of lamellipodial protrusion

Software-aided, semi-automatic cell edge tracking and analysis allowed to define 20 parameters, 18 of which are separated into groups associated with protrusion-, retraction-, dynamics/velocity- or curvature/geometry-related characteristics of the cell edge (for description of each parameter, see Table S1). The averaged and normalized values of these 20 parameters from B16-F1 wildtype *versus* Lpd KO cells are provided (for pooled data see Fig. 4 and individual clones Fig. S2). Multiple conclusions can be drawn from reviewing the 20 morphodynamic parameters in Figure 4. First, a significant reduction of *Average Advancement Velocity* (parameter #1) was observed in the absence of Lpd, confirming the results from our manual, kymograph-based analysis (Fig. 1D). Aside from parameter #1, which considers both protrusion and retraction episodes, we also dissected protrusion and retraction events individually. This analysis indicates that Lpd knockout cells display reduced values of some, but not all, protrusion-related parameters including *Effective Protrusion* (parameter #2; Figs. 4 and S2). This parameter only considers the rate of lamellipodial protrusion at the measured edge, without retraction, but averaged for the entire duration of acquisition (120 sec). Another parameter reduced in a statistically significant fashion by loss of Lpd is the *Protruding Edge Fraction* (parameter #3), which reports how many pixels on average of analysed periphery spend protruding. The reduction in parameters 2 and 3 reveals that lamellipodial protrusion activity in the absence of Lpd is compromised not only by the effective, average rate of protrusion, but also the percentage of cell periphery capable of protruding at any given time. We also found that Lpd knockout cells display reduced average velocity during individual protrusion episodes (*Protrusion Episode Velocity*, parameter #5). The difference in values obtained from this parameter, however, is subtler than for parameters #1 and #2. Furthermore, when maximal protrusion rates are displayed (*Maximal Velocity during Protrusion*, parameter #6), it becomes evident that Lpd knockout cells can transiently reach the same protrusion speed as B16-F1 controls, although this rate is not maintained as efficiently as in the presence of Lpd, explaining reductions observed in protrusion-related parameters #2-5.

**Fig. 4.**
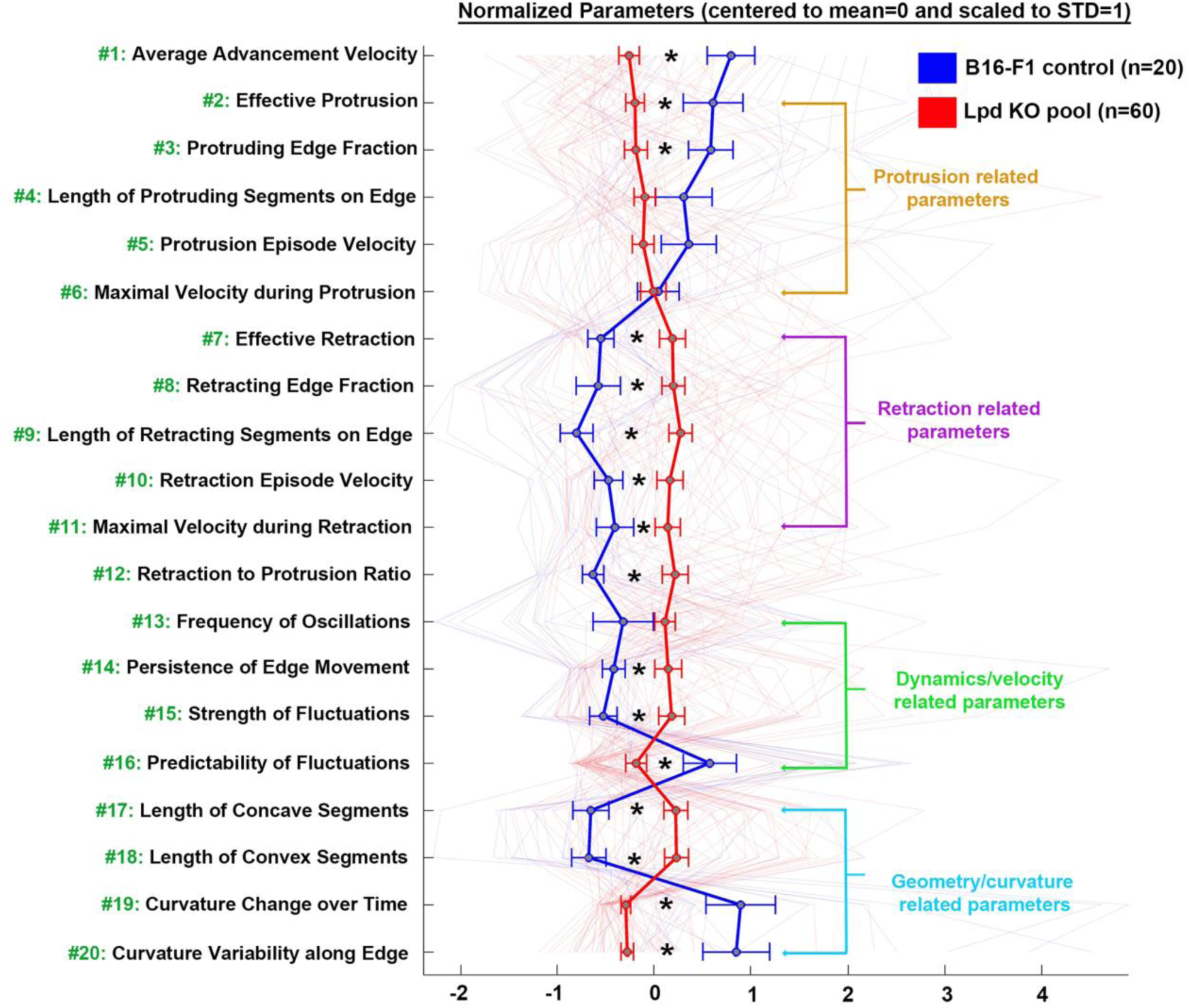
Morphodynamic analysis of lamellipodial protrusion of B16-F1 control and Lpd knockout cells. Parameters characterizing protrusion-, retraction-, velocity- or geometry-related characteristics of cell edge movements in B16-F1 wildtype and Lpd knockout cells, as derived from quantitative, morphodynamic analysis of lamellipodial protrusion. Data are represented as arithmetic means ± SEM (asterisks indicate statistically significant differences, with * indicating p<0.05). In order to facilitate the visualization of relative differences in lamellipodial morphodynamics between wildtype (marked with blue line) and Lpd knockout cells (marked with red line), 20 selected morphodynamic parameters (indicated with green numbers on y-axis) were each normalized by assigning their respective means to 0 and scaling their STD values to 1. For description of each parameter, see Table S1.

Our analysis of *Average Advancement Velocity* (parameter #1) in cells lacking Lpd reveals it is also influenced by retraction-related parameters (#7-11), as their average values are all increased in the absence of Lpd. This then contributes to the observation that the *Retraction to Protrusion Ratio* (parameter #12) is significantly increased in Lpd knockout cells as compared to controls. Furthermore, examination of the dynamics of lamellipodia reveals that relative to B16-F1 controls, Lpd KO cells, on average, tend to display more frequent membrane oscillations/fluctuations (*Frequency of Oscillations*, parameter #13) and velocity changes (*Variance of Edge Acceleration*, parameter #14), and have stronger and less predictable fluctuations (parameters #15 and #16, respectively). In spite of this variability in velocity-related features, curvature-related parameters indicated that the leading edges of Lpd knockout cells have longer concave and convex segments (parameters #17 and #18), caused by the fact that changes of curvature profile over time (*Curvature Change over Time*, parameter #19) and along the edge (*Curvature Variability along Edge*, parameter #20) are strongly reduced in the absence of Lpd. All these data suggest that Lpd also contributes to local flexibility and plasticity of cell edge protrusion, constituting a feature not previously dissected out for any protrusion regulator characterised so far.

Focusing on individual morphodynamic parameters allows drawing specific conclusions on how individual aspects of lamellipodial protrusion, retraction, dynamics or geometry differ between Lpd knockout and wildtype cells. However, analysis of individual parameters does not uncover the precise quantitative contributions of all protrusion-*versus* retraction-related parameters to the overall phenotype. Employing more sophisticated data analysis approaches can provide tools for potentially extracting such information. To this end, we applied principal component analysis (PCA), which reduces multi-dimensional data sets to two hybrid parameters with specific characteristics - principal component 1 (PC1) and principal component 2 (PC2) - using the data obtained from our morphodynamic analyses for Lpd knockout and B16-F1 control cells (Fig. 5). Static parameters such as those related to geometry/curvature were excluded in this case, allowing us to investigate features that relate to cell edge dynamics. Eleven parameters were found to have the highest contribution to PC1, the majority of which define fluctuation- or retraction-related characteristics of the cell edge (Fig. 5A). In contrast, the majority of parameters with the highest contribution to PC2 were those associated with protrusion-related characteristics of the cell edge (Fig. 5B). Thus, PC2 can be considered to define “protrusion activity”, while PC1 “retraction activity”. PC1 and PC2 contain 88% of the total variance of the data set. When displayed in a 2D-coordinate system, results revealed that high PC1 values (abscissa) almost entirely include cells of the Lpd knockout population, suggesting higher retraction activity upon loss of Lpd (Fig. 5C). In contrast, the majority of B16-F1 control cells display low or even negative values for PC1, indicating much lower retraction activity. In contrast, both B16-F1 wildtype and Lpd knockout cells seem to be equally spread throughout the PC2-axis, indicating higher heterogeneity in protrusion features within both populations. Together with the individual assessment of morphodynamic parameters described above, these results suggest that the major phenotype upon deletion of Lpd is characterised by an enhancement of different aspects of retraction activity. By considering the distribution of individual cells on the PC1/PC2 2D-coordinate system, two regimes of lamellipodial activity are apparent. The first regime, containing a mixture of Lpd knockout cells and the majority of cells in the B16-F1 control population, was characterized with negative correlation between PC1 and PC2, indicating that cells of higher protrusion activity display lower retraction activity. Interestingly, a second regime of lamellipodial protrusion, including the majority of the Lpd knockout population, featured a positive correlation between PC2 and PC1. This implies that cells with greater retraction activity also have higher protrusion activity in this regime (Fig. 5C). In order to further investigate this counterintuitive behaviour of lamellipodial dynamics, we examined the correlation between values of *Maximal Velocity during Protrusion* (parameter #6) and *Maximal Velocity during Retraction* (parameter #11), irrespective of genotype, but considering and separately displaying each of the three lamellipodial protrusion patterns, smooth, intermediate and chaotic (Fig. S3A-D). We found that a positive correlation exists between these two parameters, with cells of the chaotic phenotype displaying both the highest velocity during protrusion and the highest velocity during retraction. Only with cells of smooth phenotype, most of which harbour Lpd, we failed to detect a statistically significant correlation between protrusion and retraction velocities (Figs. 3, S3A-D). Finally, we also assessed the probability of a cell of given genotype to fall into either regime. Thus, the probability to follow regime 1 was app. 0.8 (1 representing 100%) for a given control cell, but dropped down to half if lacking Lpd, and accordingly rising from 0.2 (control) to 0.6 (KO) roughly in case of regime 2. Together, these data show that loss of Lpd can bias stronger and more frequently occurring changes in the direction of cell edge movements, coinciding with compromised lamellipodial stability.

**Fig. 5.**
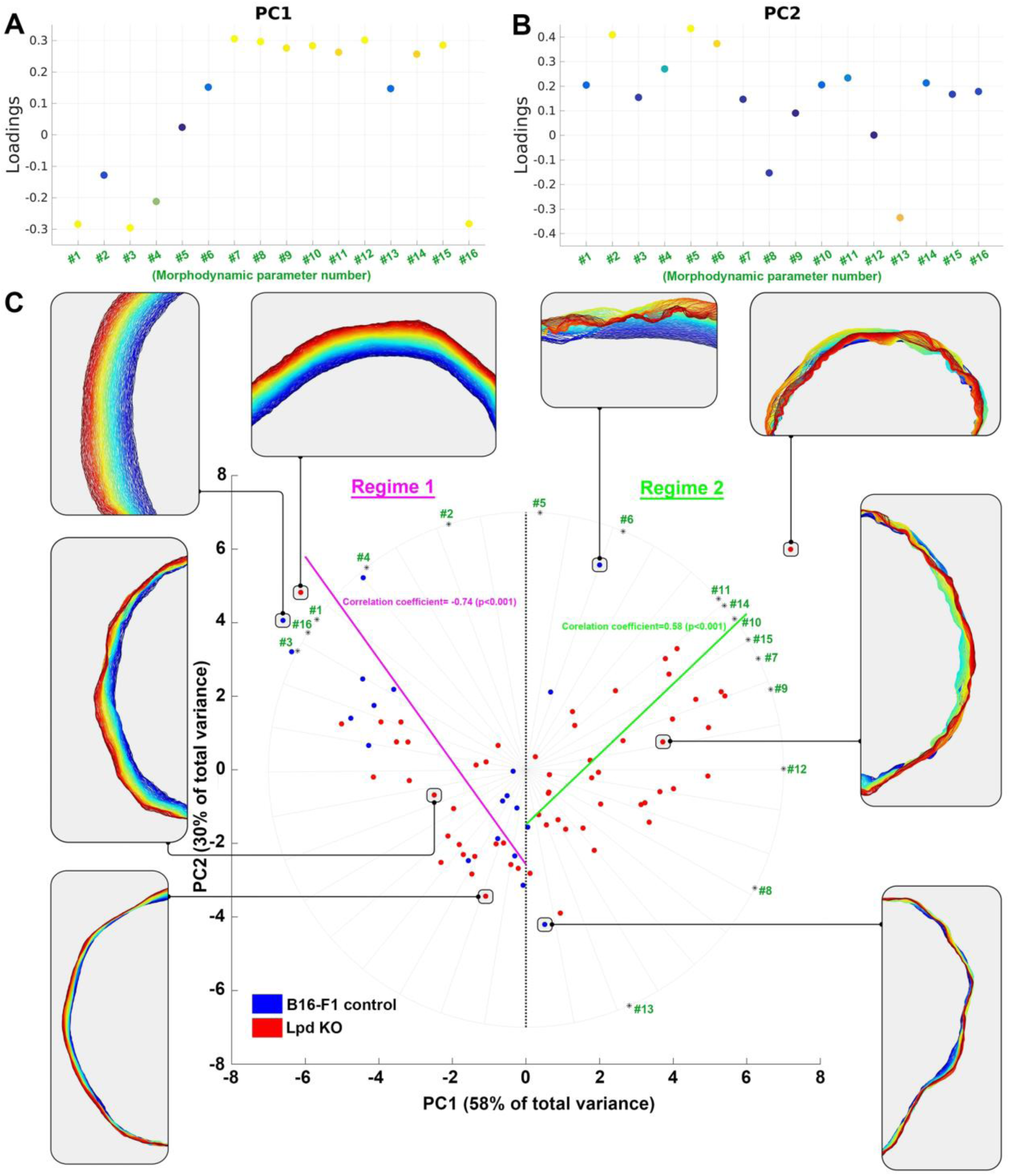
PCA analysis reveals distinct lamellipodial protrusion regimes for B16-F1 control *versus* Lpd knockout cells. (A, B) Figures display the loadings contributed by the individual parameters (derived from quantitative morphodynamic analysis) to principal components 1 and 2 (A: PC1, B: PC2). The larger the absolute value of a given loading, the stronger a given parameter contributes to respective PC. Loadings of high contribution are marked in yellow/orange. The parameter numbers correspond to those displayed in Figure 3 and Table S1, except for curvature/geometry-related parameters, which were excluded. Based on respective parameter contributions, PC1 mostly reflects retraction, and PC2 protrusion activity. (C) B16-F1 control cells (blue) and Lpd knockout cells (red) are plotted in a 2D-coordinate system with PC1 (containing 58% of total variance) and PC2 (containing 30% of total variance) on x- and y-axis, respectively. Each dot indicates a single cell, and spatiotemporal, lamellipodial contours of 8 selected cells (circles) are arranged radially around the graph, to exemplify the dynamic, lamellipodial patterns of individual, representative cells located in different sections of the PC1-PC2-plane (time-colour code as described for Fig. 2A). Cells with enhanced retraction are characterized by higher PC1 values and thus shifted towards the right of the x-axis. Cells with enhanced protrusion are characterized with higher PC2 values and shifted towards the top of the y-axis. Based on this type of data representation, two major protrusion regimes can be distinguished in our cells. Regime 1, containing most cells of the B16-F1 wildtype population and characterized by a negative correlation between PC1 and PC2 (indicated with pink line and Spearman correlation coefficient of −0.74, p<0.001). Probability of B16-F1 wildtype cells to fall into Regime1 was calculated to be 0.8 (with 1 equalling 100%), while that of Lpd KO cells 0.4. Regime 2, comprised mostly of cells of the Lpd knockout population and characterized by a positive correlation between PC1 and PC2 (indicated with light green line and Spearman correlation coefficient of 0.58, p<0.001). Probability of B16-F1 wildtype cells to fall into Regime 2 was calculated to be 0.2, while that of Lpd KO cells 0.6. Dominance of a specific parameter in the characterization of the morphodynamics of a cell corresponds to a parameter-specific direction in the PC1-PC2-plane (green numbers). The morphodynamic characterization of cells positioned on a line from the origin to a specific green number is dominated by the parameter with this number.

### Lpd knockout does not alter the lamellipodial actin network

Our analysis reveals that Lpd deletion reduces net lamellipodial protrusion, associated with reduced persistency of protrusion caused by enhanced retraction and oscillation events. Given this, we investigated whether loss of Lpd results in general morphological or dynamic defects in lamellipodial actin networks. To our surprise, there were no obvious morphological defects or significant differences in intensity or width of lamellipodial actin in Lpd knockout cells, as compared to B16-F1 wildtype cells (Fig. 6A, B). Furthermore, loss of Lpd does not impact on the rate of actin network polymerisation and the intensity of Arp2/3 complex in lamellipodia (Fig. S3E-H). These observations indicate that reduced lamellipodial protrusion and cell migration rates upon deletion of Lpd are not caused by defects in lamellipodial actin network formation and Arp2/3 complex incorporation.

**Fig. 6.**
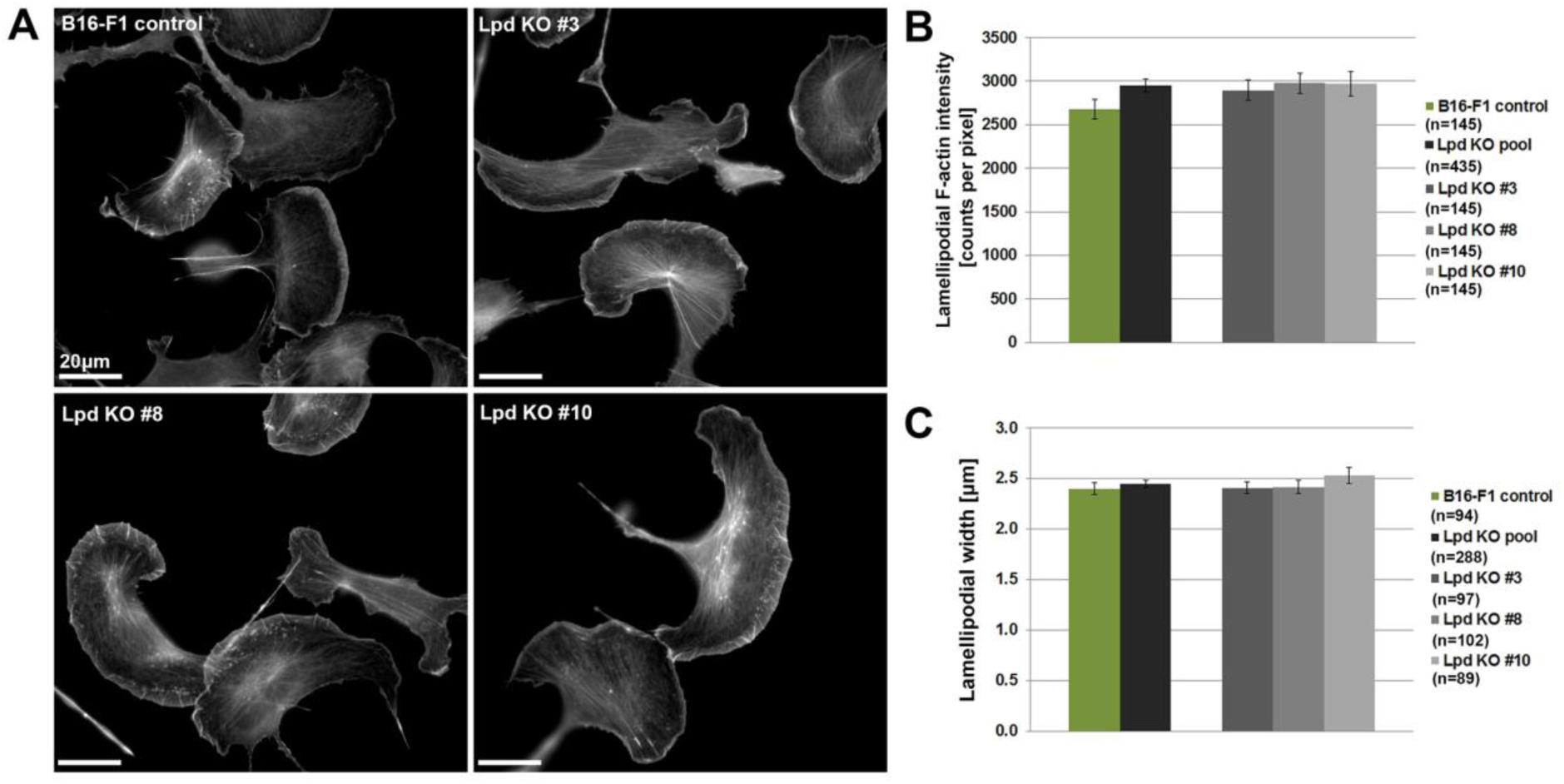
Lpd KO cells lack discernible defects in lamellipodial actin networks. (A) B16-F1 cells of indicated genotype migrating on laminin, and fixed and stained for F-actin with phalloidin. (B) Graphs show quantitation of lamellipodial F-actin intensity and (C) lamellipodial width of individual Lpd KO clones and the pooled population compared to controls. n=number of cells, and data are represented as arithmetic means ± SEM. Statistics revealed no significant differences between any pair of experimental groups (not shown).

### Lpd stabilizes VASP, but does not recruit VASP or Abi to lamellipodia

Lpd has previously been implicated in the recruitment and/or stabilization of Ena/VASP proteins at the lamellipodium edge or their tethering to growing actin filaments (Bae et al., 2010; Hansen and Mullins, 2015; Krause and Gautreau, 2014; Krause et al., 2004; Michael et al., 2010; Pula and Krause, 2008). We thus speculated that potential alterations in VASP dynamics and/or regulation might be responsible for the defects in protrusion caused by the loss of Lpd. However, there was no discernible difference in VASP localization or intensity at the very front of lamellipodia between Lpd knockout and B16-F1 wildtype cells (Fig. 7A, B). Similar results were obtained for Mena (Fig. S3I, J). Immunofluorescence analysis shows that VASP and Mena accumulate at lamellipodial edges in the absence of Lpd, but does not address whether Lpd affects their kinetics. Using FRAP-based approaches, we found that the half-time of recovery of VASP but not the overall extent is reduced in the absence of Lpd as compared to B16-F1 controls (Fig. 7C, D). These data suggest that Lpd contributes to the stabilization of Ena/VASP proteins at lamellipodial edges, in spite of being dispensable for their recruitment.

**Fig. 7.**
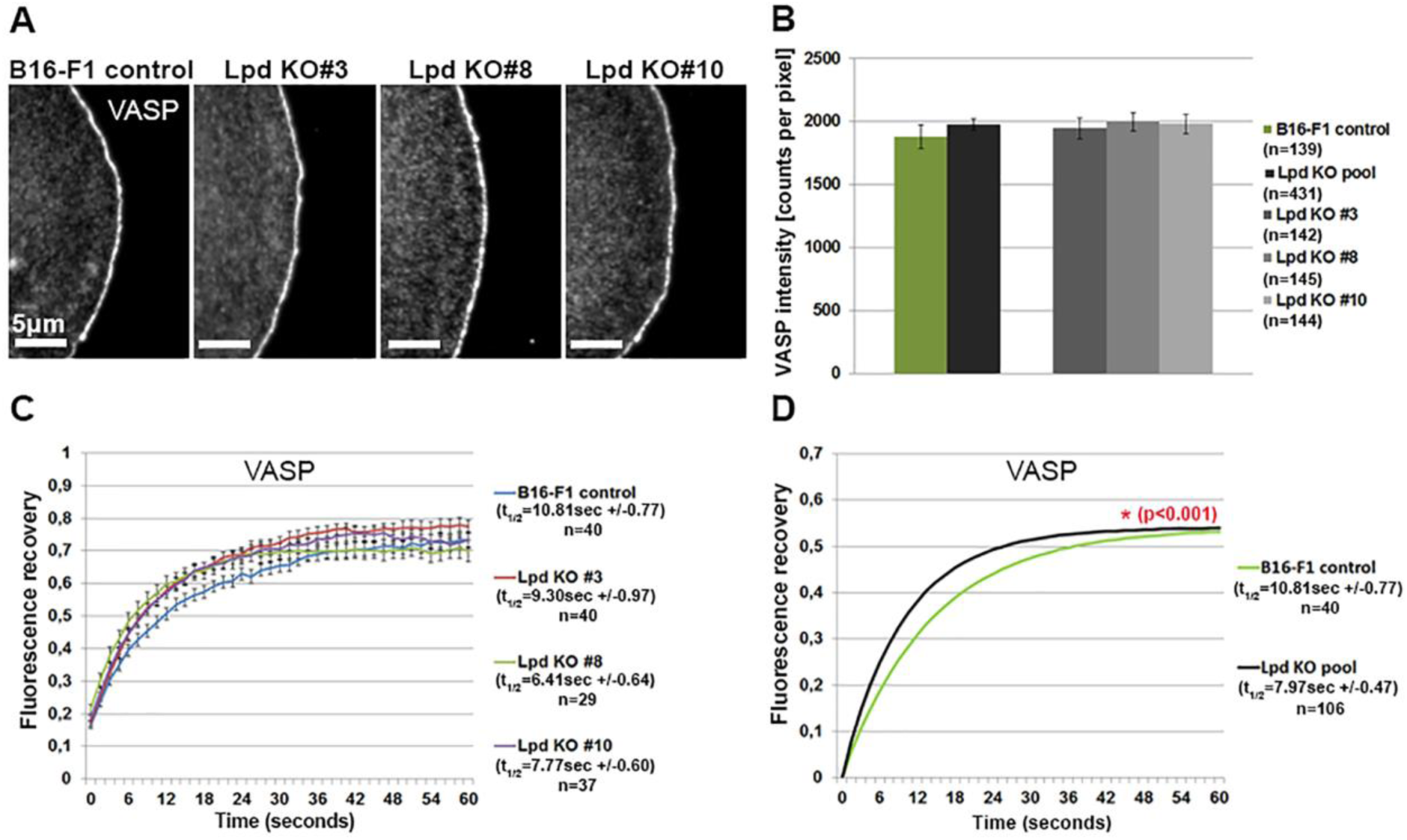
Analysis of VASP intensity and turnover at lamellipodia edges in the presence and absence of Lpd. (A) Immunofluorescence analysis showing the localization of VASP at lamellipodia edges in both B16-F1 control and Lpd knockout cells. (B) Quantification of VASP intensities at lamellipodia edges in B16-F1 wildtype and Lpd knockout cells, displayed individually and as pooled population. n = number of cells, and data are arithmetic means ± SEM. (C) The graph shows the recovery curves of EGFP-tagged VASP at the leading edge after bleaching in B16-F1 cells and individual Lpd knockout clones, as indicated. Half-times of recovery (in seconds) obtained from curve fitting are displayed on the right. Each time point after the bleach is represented as arithmetic mean ± SEM. (D) Fluorescent recovery curve fits of lamellipodial FRAP data of B16-F1 wildtype and pooled Lpd KO cells as well as derived t_1/2_-values, as indicated on the right. n = number of FRAP events analysed. A significant statistical difference in t_1/2_-values between B16-F1 wildtype and pooled Lpd KO cells is indicated with red colour (p<0.001).

Lpd is thought to promote cell migration via its interaction with the WAVE Regulatory Complex (WRC) during lamellipodium protrusion (Law et al., 2013). Immunofluorescence analysis of the WRC subunit Abl interacting protein (Abi) reveals that WRC accumulation at lamellipodia edges in the absence of Lpd is increased rather than decreased compared to control B16-F1 cells (Fig. S4A, B). However, EGFP-Abi (WRC) dynamics at the leading edge was identical in both cell types (supplementary Fig. S4C, D). This suggests that the presence of Lpd antagonises WRC recruitment to the lamellipodium edge by a hitherto unknown mechanism. Immunoblotting confirmed there were no changes in expression levels of VASP as well as the WRC subunits Abi, Nap1 and WAVE2 in Lpd knockout clones compared to B16-F1 wildtype (Figs. S4E-H). These results exclude that the relatively modest phenotypes we see are due to compensatory changes in expression levels of respective components.

### Lpd loss reduces nascent adhesion formation underneath lamellipodia

Since cells lacking Lpd are characterized by fluctuating and rapidly retracting lamellipodia, we hypothesized that lack of stability of these structures might be caused by defects in adhesion to the substratum. In order to test this hypothesis, we expressed EGFP-paxillin in Lpd knockout and B16-F1 control cells, and performed live cell imaging to quantify the number of nascent adhesions during lamellipodia formation (Fig. 8A). We distinguished between front and back halves of the lamellipodium and found that nascent adhesion distributions strongly depended on respective lamellipodial protrusion class in both cell types (Fig. 8A and Movie 4). Taking all protrusion classes together, there was a clear reduction in the number of new adhesions in the absence of Lpd (Fig. 8B, left panel). Subcategorization showed that the number of new adhesions was not significantly reduced in the absence of Lpd in cells with smooth or chaotic protrusions (Fig, 8B, right panel). In contrast, the intermediate class of protrusions had a statistically significant reduction in adhesion number if Lpd was absent. This combined with the fact that the chaotic phenotype (with much lower adhesion numbers) increased in frequency in Lpd KO cells (Fig. 3D) explains the overall reduction of new adhesions in the absence of Lpd (Fig. 8B).

**Fig. 8.**
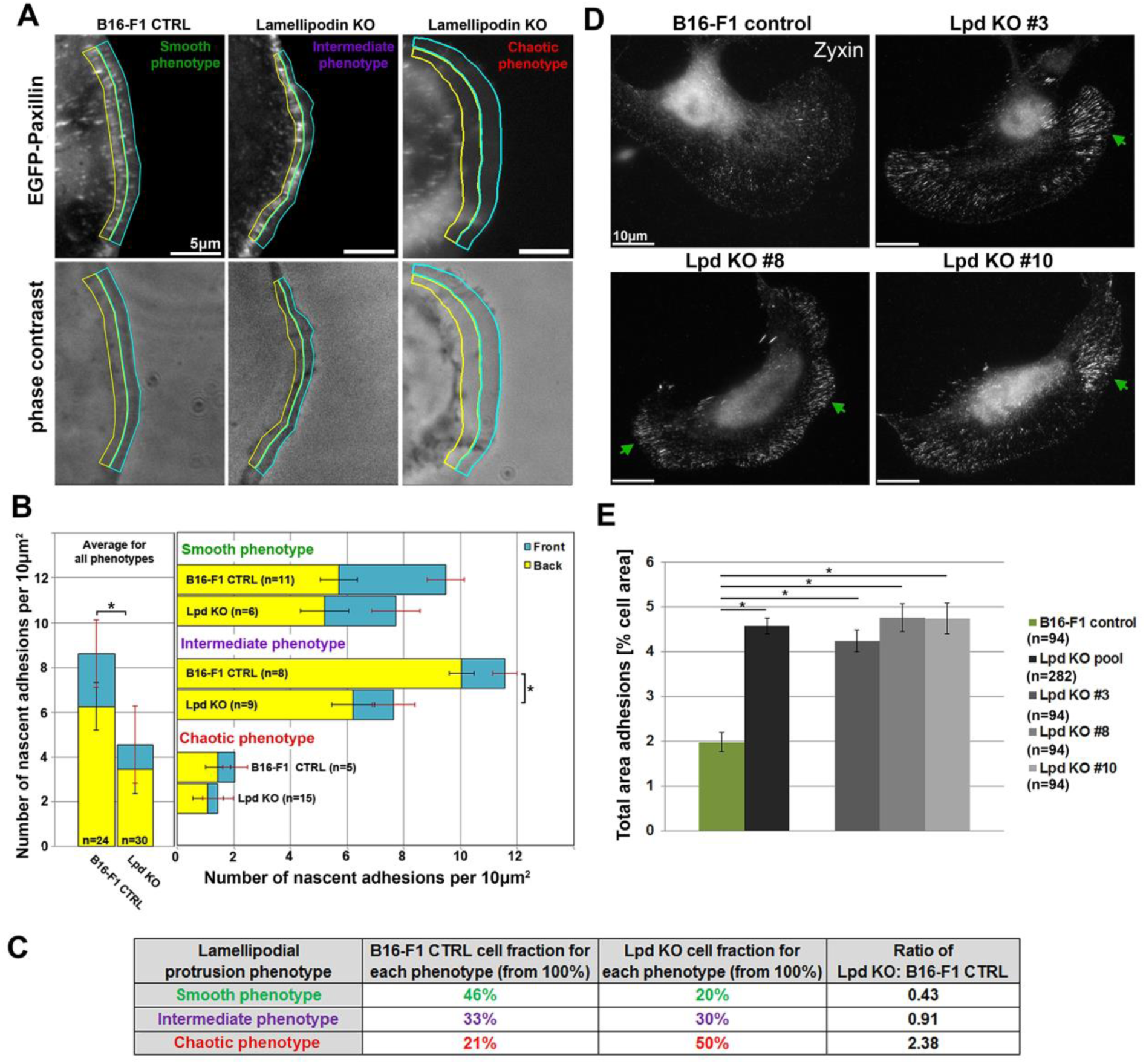
Lpd deletion compromises nascent adhesion and promotes mature, focal adhesion formation. (A) Representative, EGFP-paxillin and phase contrast images of protrusive regions of B16-F1 wildtype or Lpd knockout cells, with the front halves of their lamellipodia encircled with cyan, and the back halves with yellow. Images display distributions of nascent adhesions in the three distinct lamellipodial protrusion phenotypes defined in Figure 3. (B) Quantification of nascent adhesion numbers in lamellipodia of each protrusion phenotype (smooth, intermediate or chaotic), as well as averaged for all phenotypes and displayed for both genotypes (B16-F1 wildtype and Lpd KO) separately (left). Quantifications are also separated for front and back lamellipodial regions, again highlighted as cyan and yellow bars, respectively. (C) Table indicating cell fractions belonging to each protrusion phenotype (smooth, intermediate or chaotic) for both control and Lpd KO cell populations upon EGFP-paxillin expression. The Lpd KO population corresponds to data from Lpd KO#3. The ratio of Lpd KO over B16-F1 wildtype cells for each protrusion phenotype is shown in the fourth column on the right, again confirming significantly reduced cell numbers among Lpd knockout cells as compared to controls harbouring the smooth lamellipodial phenotype, and *vice versa* for the chaotic phenotype. (D) B16-F1 wildtype or Lpd knockout clones were seeded onto laminin and mature focal adhesions stained with Zyxin-reactive antibody. Green arrows point at typical clusters of focal adhesions more frequently observed in Lpd knockout clones. (E) Quantitation of average focal adhesion area (relative to cell area) as determined in distinct cell types, as indicated, demonstrating that Lpd removal enhances the area occupied by focal adhesions. All data are arithmetic means ± SEM, with asterisks above bar charts indicating statistically significant differences between designated groups (* = p<0.05), n equals numbers of cells analysed.

Although cells expressing EGFP-paxillin displayed an increase of the smooth, i.e. most efficiently protruding phenotype at the apparent expense of the intermediate phenotype in both genotypes (Fig. 8C, compare with 3D), our analysis suggests that Lpd deficiency reduces the average probability of nascent adhesions to be formed below newly protruding lamellipodia. A closer look at the spatial distribution of adhesion formation reveals approximately a third of new adhesions form at the front part of smoothly protruding lamellipodia (Fig. 8B and Movie 4). This is consistent with new adhesions being continuously formed close to the front, with a subset being stabilized and maturing into focal adhesions, as the lamellipodium persistently moves forward. In contrast, despite having a higher total number of adhesions, cells with intermediate protrusions have significantly lower ratio of adhesion numbers in front over back parts of the lamellipodium (Fig. 8B). Moreover, adhesions appeared to accumulate at the lamellipodium-lamella border, with lower numbers of new adhesions at the front. Finally, cells with chaotic lamellipodial phenotype had very low numbers of nascent adhesions compared to the other two classes, which probably accounts for their lack of stability and frequent cycles of protrusion and retraction or collapse.

We thus conclude that Lpd helps stabilize and/or maintain nascent adhesions, which must occur indirectly since Lpd does not apparently localize to these adhesions (Fig. S1C, Movie 5). One of the earliest regulators of actin dynamics implicated in nascent adhesion formation constitutes the Rho-family GTPase Rac, which is also essential for lamellipodia protrusion and promotes the formation of Rac-induced focal complexes (Nobes and Hall, 1995; Rottner et al., 1999b; Schaks et al., 2018; Steffen et al., 2013). We thus explored to what extent the reduction of nascent adhesion upon Lpd KO occurs through potentially compromised Rac signalling. Firstly, Lpd KO clones are completely unchanged as compared to controls in their respective activities of endogenous Rac GTPases (Fig. S5A). Secondly, although ectopic Rac1 (wildtype) expression increases the number of EGFP-paxillin-containing, nascent adhesions in Lpd KO cells, the situation is complex, as the opposite trend was seen upon Rac-transfection in control cells (Fig. S5B). Likewise, EGFP-Rac expression and subsequent analysis of protrusion parameters revealed that a sole increase of wildtype Rac levels increases *Average Advancement Velocity* (parameter #1) only in Lpd KOs (and not in controls, Fig. S5C), and this is due to counteracting retraction and not by increasing protrusion (compare e.g. parameters #5 and #12 in S5D and S5E). Comparison of all parameters showed that retraction-related parameters (#7-#12) are reduced upon Rac expression in Lpd KO cells. Nevertheless, these data and the variety of changes observed upon Rac1 expression in both control and Lpd KO cells clearly illustrates that changes in Rac level or activity are not causative of phenotypes in protrusion and nascent adhesion observed upon Lpd removal (Fig. S6). Consistent with aforementioned role of Lpd in stabilization and/or maintenance of nascent adhesions, Lpd loss also strongly shifted the balance between nascent and mature adhesions towards the latter, as Lpd KOs display an increased area of more mature adhesions containing zyxin (Fig. 8D, E). This shift may explain the reduced efficiency of early, integrin-dependent cell spreading, in particular observed on fibronectin, but also, albeit less prominently, on laminin (Fig. S4I, J). Taken together, our data establish that loss of Lpd leads to a Rac signalling-independent defect in nascent adhesion formation accompanied by their increased maturation, likely contributing to the destabilization of lamellipodial protrusions.

### Lpd regulates the number but not speed of Vaccinia-induced actin tails

Vaccinia virus promotes its actin-dependent cell-to-cell spread at the plasma membrane by activating the Arp2/3 complex in a Nck, WIP and N-WASP dependent fashion (Frischknecht et al., 1999b; Moreau et al., 2000; Welch and Way, 2013). Curiously, and like lamellipodia protrusion, this pathway also involves Rac signalling at the plasma membrane (Alvarez and Agaisse, 2013). Furthermore, as Vaccinia motility and lamellipodium protrusion share various features of actin-based motility, such as dependence on Arp2/3 complex activation at the plasma membrane as well as recruitment of Lpd (Krause et al., 2004) and VASP (Frischknecht et al., 1999a; Grosse et al., 2003), we investigated whether Lpd depletion affects motility of the virus. Consistent with previous immunofluorescence analysis (Krause et al., 2004), we first confirmed that both EGFP-tagged and endogenous Lpd are recruited to Vaccinia-induced actin tails in HeLa cells (Fig. S7A). Indeed, RNAi-mediated knockdown of Lpd in these cells reduced the number of Vaccinia-induced actin tails (Fig. S8A, B). Effective Lpd knockdown could be documented both by immunoblotting of bulk cell populations (Fig. S8A) and immunolabelling at the single cell level (Fig. S7B). However, the length of actin tails and speed of viral movement, the latter of which is powered by actin polymerization, were unchanged upon Lpd knockdown (Fig. S8B). Defects on Vaccinia tail formation frequency occurred in spite of equal numbers of extracellular virus, so derived from effects on tail initiation and/or maintenance (Fig. S7C). Similar results were obtained upon tamoxifen treatment-mediated Lpd gene removal (Lpd KO) in MEFs originating from conditional Lpd (Lpd *fl/fl*) knockout mice (Fig. S8C, D; Law et al., 2013). Both tail formation frequency and tail length were reduced in the absence of Lpd in these fibroblasts, but viral speed as direct readout of actin assembly was again unchanged (Fig. S8D). This observation was fully consistent with recruitment in the absence of Lpd of Nck and N-WASP, the latter of which operates as essential Arp2/3 complex activator in the Vaccinia system and in analogy to WRC in lamellipodia, as well as of the Lpd-interactor VASP (Fig. S7D). Taken together, these experiments suggest that Lpd contributes to formation frequency and maintenance of specific actin structures at the plasma membrane, such as lamellipodia and Vaccinia-induced actin tails, but its removal does not eliminate these structures nor compromise the efficiency of actin filament assembly in them.

## DISCUSSION

Here, we explored the mechanistic function of lamellipodin (Lpd) in migrating B16-F1 cells using CRISPR/Cas9-mediated gene disruption. In agreement with previous, RNAi-mediated knockdown and conditional knockout of Lpd in MEFs (Krause et al., 2004; Law et al., 2013), our data confirm that Lpd is a positive regulator of lamellipodial protrusion and cell migration on 2D substrates. Importantly though, it is not obligatory for lamellipodia formation in all cell types and conditions, consistent, for instance, with reduction but not loss of lamellipodia formation upon Lpd knockdown in MTLn3 breast cancer cells (Carmona et al., 2016). However, we cannot exclude compensatory mechanisms occurring during permanent or more short-term CRISPR-treatment in our model systems, since tamoxifen-induced, conditional Lpd knockout MEFs had previously shown impaired lamellipodia formation and stronger migration phenotypes than those observed here (Law et al., 2013). Nevertheless, loss of Lpd does not phenocopy loss of WRC (see e.g. Schaks et al., 2018). This is also consistent with the observation that Lpd knockout in mice is compatible at least with embryonic development and life until early after birth (Law et al., 2013). However, this contrasts with mice lacking obligatory lamellipodial regulators, such as WRC subunits (Rakeman and Anderson, 2006; Yamazaki et al., 2003) or Rac1 (Sugihara et al., 1998), which display early embryonic lethality. The approach of growing permanently deleted cell lines allowed us to precisely explore mechanistic changes in lamellipodia formation accompanied by Lpd loss of function, while key features of our phenotypes could also be confirmed upon more short-term, CRISPR/Cas9-mediated gene disruption. Migration of B16-F1 melanoma cells on laminin is characterized by distinct protrusion phenotypes, varying from “smooth” to “chaotic” as two extremes as well as an “intermediate” phenotype, the relative frequencies of which change upon Lpd knockout. By means of semi-automated cell edge tracking combined with quantitative, morphodynamic analysis, we found that defects in lamellipodial persistence in Lpd knockout cells are associated with an increase in the retraction and oscillation features of their cell edges. Surprisingly, however, Lpd knockout cells are capable of reaching lamellipodial protrusion velocities similar to those of wildtype cells. These observations agree with the lack of detectable lamellipodial actin network defects, including F-actin densities, lamellipodial Arp2/3 complex incorporation or actin polymerisation rates. Due to the coincident increase rather than decrease of accumulation of its interaction partner Abi-1, subunit of the lamellipodial Arp2/3 complex activator WRC, we conclude the physical interaction between Lpd and Abi to have regulatory functions instead of contributing to WRC recruitment and/or accumulation. As opposed to the WRC-subunit Abi, Lpd can increase the dwell time at the lamellipodium edge of the Ena/VASP family member VASP, which may also affect the activity of the latter on growing actin filaments, as suggested previously (Hansen and Mullins, 2015). However, such changes are likely too subtle for detection in the context of the multitude of factors potentially regulating lamellipodial actin polymerization and protrusion.

When comparing the impact of Lpd on the two different types of actin-based protrusion at the plasma membrane studied here, lamellipodia and Vaccinia virus-induced actin tails, we found two major commonalities. First, little effect on upstream regulation and incorporation of the major actin assembly factor in these protrusions, the Arp2/3 complex, and second, a peculiar contribution in both cases to formation frequency and stability of these structures. More specifically, the lamellipodia in cells lacking Lpd had problems with efficiently advancing forward, at least on average, and displayed a higher likelihood to chaotically collapse or retract. Thus, an increased tendency to retract into the cortex might also be the cause for reduced Vaccinia tail numbers, if indeed analogous. If translating this into lamellipodial activity, an increased probability of continuous, smooth protrusion, as observed in the presence of Lpd, will clearly correlate with the likelihood of depositing material for nascent adhesion formation. And indeed, although not essential, the presence of Rac and thus protruding lamellipodia is known to clearly promote nascent adhesion formation by increasing the mobile fractions of various actin binding proteins present in these structures, including paxillin, zyxin and VASP (Steffen et al., 2013). Consistently, numbers of nascent adhesions in lamellipodia of Lpd knockout cells were found, on average, reduced relative to those in B16-F1 controls, and those of mature focal adhesions increased. These data suggest a shift in activities from protrusive, Rac-dependent behaviour towards more contractile, Rho-dependent activity in the absence of Lpd. However, this does not mean that we can explain defects observed upon Lpd removal by sole reduction of Rac signalling, or rescue them by increased Rac activity. In fact, experiments revealed that Lpd removal did not impact on Rac activity, and upregulation of cellular Rac dose, in spite of creating a condition for improvement of retraction-related phenotypes and slight increase of nascent adhesion numbers in Lpd KO cells, did not restore the complexity of differences in protrusion behaviour between control and Lpd KOs.

Aside from this, apparent defects in adhesion regulation may also be linked to the various interactions of Lpd with established adhesion components such as talin or integrins (Lagarrigue et al., 2015; Lee et al., 2009; Watanabe et al., 2008). Moreover, a potential involvement of Rac in Lpd-dependent regulation of nascent adhesions may derive from defective transduction of extracellular matrix engagement through FAK and Cas signalling (Bae et al., 2014), but mechanistic details remain to be determined. Whatever mechanism, Lpd loss of function clearly increases the likeliness of retraction and rearward-folding of lamellipodia.

Repeated cycles of protrusion and retraction were previously reported for a variety of cell types and with a broad range of periods and amplitudes (Burnette et al., 2011; Enculescu et al., 2010; Gholami et al., 2008; Giannone et al., 2004; Giannone et al., 2007; Koestler et al., 2008; Machacek and Danuser, 2006; Zimmermann and Falcke, 2014). The chaotic behaviour that we find in Lpd KOs shows similarities to spreading fibroblasts exhibiting so called periodic contractions, which coincide with brief interruptions of protrusion accompanied by integrin and adhesion clusters formation close to the cell edge (Giannone et al., 2004; Giannone et al., 2007). Although protrusion and retraction events are less clearly separated in B16-F1 lamellipodia protruding on laminin, Lpd appears to interfere with the coordination of these activities. The precise changes observed upon Lpd removal concern multiple parameters, either reflecting decreased protrusion and increased retraction or cell edge activities to become less predictable, less robust, with lower plasticity and overall more chaotic. The combination of all these phenotypic parameters reduces the efficiency of protrusion and nascent adhesion formation and turnover for productive forward advancement of the lamellipodium, and ultimately the efficiency of cell migration. While confirming this particular aspect of previously published literature and at the same time specifying the phenotypic changes caused by Lpd removal, the precise mechanistic reasons for these changes remain to be elucidated.

## MATERIALS AND METHODS

### Cell culture, transfection conditions, RNAi and induced Lpd knockout

B16-F1 mouse melanoma cells (purchased from American Type Culture Collection, Manassas, VA) were cultivated according to standard tissue culture conditions. Cells were grown at 37°C / 7.5% CO_2_ in DMEM medium containing 4.5 g/l glucose (Life Technologies, Thermo Fisher Scientific, Germany) and 10% fetal calf serum (FCS, PAA Laboratories, Linz, Austria), 2 mM glutamine (Life Technologies), and 1% penicillin-streptomycin (Life Technologies). For transfections of cells with DNA vectors described below, JetPei transfection reagent (Polyplus Transfection, Illkirch, France) was used according to manufacturer’s instructions. For microscopy experiments, B16-F1 cells were plated onto glass coverslips pre-coated for 1 hour at room temperature with 25 μg/ml laminin (L-2020; Sigma-Aldrich) in 50 mM Tris, pH 7.4, and 150 mM NaCl or 25 μg/ml fibronectin (Roche) in PBS.

Rat2 fibroblasts and NIH-3T3 mouse fibroblasts were grown at 37°C / 7.5% CO_2_ in DMEM medium containing 4.5 g/l glucose and 10% FBS (054M3396, Sigma-Aldrich, Germany), 2 mM glutamine, 1 mM sodium pyruvate (Life Technologies), 1% non-essential amino acids (Life Technologies) and 1% penicillin-streptomycin.

Hela cells were maintained in DMEM supplemented with 10% FBS and antibiotics (P/S). To perform shRNA-mediated knock-down, HeLa cells were transfected with Lpd-specific or scramble shRNA as control using Fugene HD (Roche) transfection reagent, as described (Carmona et al., 2016). Following transfections, cells were incubated for 24 hours before undergoing 48-72 hours of selection with 2 μg/ml puromycin, and then subjected to viral infections and analyses as described below.

Tamoxifen (4-OHT)-inducible mouse embryonic fibroblasts (Lpd fl/fl-CreER MEFs) from conditional Lpd KO mice were generated and grown as described (Law et al., 2013). To prepare KO MEFs prior to infections, Lpd fl/fl-CreER MEFs were incubated in medium containing 0.6 µM Tamoxifen (Sigma) for 2 days, followed by 6 days incubation with 0.3 µM Tamoxifen. All cell lines in our lab are routinely authenticated for induced gene modifications, both by ourselves and local authorities, and routinely screened for mycoplasma contaminations at regular frequency.

### Generation of B16-F1 Lpd knockout cell lines

Generation of B16-F1 mouse melanoma Lpd knockout clones was performed analogous to FMNL2/3-KO clones described previously (Kage et al., 2017). CRISPR/Cas9 guide design was performed using a publicly available CRISPR design tool (back then http://crispr.mit.edu), with the DNA sequence of exon 5 of Lamellipodin/Raph1 being used (base pairs 937-1014 of cDNA of NM_001045513.3). A guide with the highest available aggregate score of 75/100 was selected, targeting the following genomic DNA sequence: 5’-TGAGAAGATCCGAGTTGCTC-3’. Forward and reverse guide oligo sequences, respectively 5’-CACCGTGAGAAGATCCGAGTTGCTC-3’ and 5’-AAACGAGCAACTCGGATCTTCTCAC-3’ were annealed for 4 min at 95°C, followed by 10 min incubation at 70°C and gradual cooling at room temperature in a buffer containing 100 mM KAc, 2 mM MgAc, 30 mM HEPES-KOH (pH=4.7). Annealed sequences were cloned into an expression vector pSpCas9(BB)-2A-Puro(px459) (Addgene plasmid ID:48138) using BbsI restriction enzyme. Successful cloning was confirmed by sequencing with primer 5’-AGGCTGTTAGAGAGATAATTGG-3’. For generation of Lpd knockout cell clones, B16-F1 cells were transfected with the cloned plasmid encoding puromycin resistance and the CRISPR guide sequence targeting Lpd. Transfected cells were cultured for 4 days with B16-F1 culture medium containing 2.5 mg/ml puromycin. Single colonies were isolated and expanded until confluent. Confirmation of genetic knockout was validated by both western blot and DNA sequencing of genomic DNA. For gDNA extraction, B16-F1 cells were pelleted and incubated at 55°C overnight in lysis buffer (100 mM Tris pH 8.5, 5 mM EDTA, 0.2% SDS, 200 mM NaCl) containing 40 µg/ml proteinase K. Standard phenol/chloroform precipitation was performed for extraction of nucleic acids. Genomic DNA sequence of 364bp, flanking the Lpd target sequence, was amplified by Phusion High-Fidelity Polymerase (New England Biolabs) with 5’-GAACGGGCCATTTTAAAATTGTGC-3’ and 5’-AGACATTAGGAAGAATACAGTTTTACC-3’ as forward and reverse primers, respectively. Amplified sequence was purified with NucleoSpin Gel and PCR clean-up kit according to the manufacturer’s instructions (Macherey&Nagel), cloned into a Zero Blunt TOPO vector using Zero Blunt TOPO Cloning Kit (Invitrogen) and transformed into competent bacteria. Single bacterial colonies were isolated and inoculated, plasmid DNA purified using NucleoSpin Plasmid kit (Macherey&Nagel) and sequenced by MWG-Biotech (Ebersberg, Germany) using sequencing primer 5’-CAGGAAACAGCTATGAC-3’. Clones with frameshift mutations on all alleles causing stop codons downstream of the target site were selected for further characterization.

### CRISPR/Cas9 treatment of Rat2 fibroblasts

Lpd gene disruption in Rat2 fibroblasts as shown in Fig. 2 was as described for B16-F1 cells above, except that cells were analysed after transient gene disruption as mixed cell populations in this case. Transfections with respective CRISPR/Cas9 construct (with specific target sequence being identical for murine and rat Lpd-sequences) were followed by 4 days of puroymycin selection and 1-2 days of recovery and growth to near-confluence of surviving cells for further processing for experiments, i.e. replating onto plastic dishes for random cell migration experiments or onto coverslips for cell stainings. Thus, cells were analysed roughly 7 days after transfections with CRISPR/Cas9 construct.

### Live-cell imaging equipment and conditions

All live-imaging experiments were performed with either inverted Axio Observer (Carl Zeiss, Jena, Germany) equipped with an automated stage, a DG4 light source (Sutter Instrument, Novato, CA) for epifluorescence illumination, a VIS-LED for phase contrast imaging, an acousto-optic tunable filter (AOTF)-controlled 405 nm diode laser for FRAP and a CoolSnap-HQ2 camera (Photometrics, Tucson, AZ), driven by VisiView software (Visitron Systems, Puchheim, Germany) or an inverted microscope (Axiovert 100 TV; Carl Zeiss), equipped with an HXP 120 lamp for epifluorescence illumination, a halogen lamp for phase-contrast imaging, a CoolSnap-HQ2 camera and electronic shutters driven by MetaMorph software (Molecular Devices, Sunnyvale, CA) for image acquisition. Cells, seeded on cover glasses, were mounted on an open heating chamber (Warner instruments, Hemden, CT) linked to a heating controller (TC-324 B) maintaining a constant temperature of 37°C. During high-magnification, live-imaging experiments, B16-F1 cells were incubated in microscopy medium (Ham’s F-12 4-(2-hydroxyethyl)-1-piperazineethanesulfonic acid–buffered medium; Sigma-Aldrich) including 10% FCS (PAA Laboratories), 2 mM glutamine, and 1% penicillin-streptomycin (both purchased from Life Technologies).

### Lamellipodium protrusion rate measurements by kymograph-based analysis

Manual determination of average lamellipodium protrusion rate was performed as previously described (Dimchev et al., 2017). Kymographs were obtained using Metamorph from time-lapse phase contrast movies of either B16-F1 wildtype cells, of Lpd knockout clones or of both overexpressing EGFP-tagged Lpd. Experiments were performed with either 63x/1.4 NA apochromatic or 100x/1.4 NA Plan apochromatic oil objectives. Average values of lamellipodial protrusion rates were displayed as μm/min.

### Random migration assays

Random migration rates of B16-F1 and Rat2 wild type and Lpd knockout cells, seeded on 25 μg/ml laminin pre-coated μ-slide four-well glass bottom microscopy chambers (Ibidi GmbH, Martinsried, Germany), were determined using phase-contrast time-lapse movies taken with a 10x/0.15NA Plan Neofluar objective over a period of 10hrs, and time interval of 10min. Microscopy chambers were mounted on a stage equipped with an incubator maintaining a constant temperature of 37°C and 7.5% CO_2._ Analysis was performed on ImageJ using the Manual tracking plugin and *Chemotaxis and Migration Tool* by Ibidi. Average values of random migration rates were displayed as μm/min.

### DNA constructs

EGFP-actin and EGFP-CAAX (pEGFP-F, farnesylated) was purchased from Clontech (Mountain View, USA), and remaining constructs published previously as follows: EGFP-Lpd (Krause et al., 2004), EGFP-Abi1 (Innocenti et al., 2005), EGFP-tagged VASP and paxillin (Rottner et al., 2001), and the Prel/RIAM expression construct, termed mmPrel1-CMV-Sport6 (Jenzora et al., 2005). pRK5-myc-Rac1 (wt) co-expressed with EGFP-paxillin for nascent adhesion formation analysis in Fig. S5B was generated by site-directed mutagenesis of pRK5-myc-Rac1-Q61L, kindly provided by Dr. Laura Machesky (Beatson Institute, Glasgow, UK). EGFP-Rac1(wt) used for the generation of data shown in Figs. S5C-E was generated by fusing human Rac1 cDNA into EGFP-C1 (Clontech) and kindly provided by Dr. Wolfgang Kranewitter (Hospital Barmherzige Schwestern, Linz, Austria).

### Immunoblotting and antibodies

For confirmation of CRISPR/Cas9-mediated gene disruption by Immunoblotting, protein lysates were prepared in a buffer containing 50 mM Tris (pH 7.5), 150 mM NaCl, 1 mM EDTA, 1% Triton X-100, supplemented with a Complete mini EDTA-free Protease Inhibitor Cocktail pill (Roche). For obtaining total protein lysates, cells were lysed in Laemmli sample buffer (60 mM Tris-Cl pH 6.8, 2% SDS, 10% glycerol, 5% β-mercaptoethanol, 0.01% bromophenol blue), syringed multiple times and incubated for 20 min with Benzonase nuclease (0.5 μl per volume of 30 μl) at 37°C in order to reduce solution viscosity caused by genomic DNA. Protein concentrations were quantified using the Pierce BCA Protein Assay Kit (Thermo Fisher Scientific). All lysates were brought to a final amount of 30μg, and boiled for 10 min at 95°C before being loaded on SDS-PAGE gels. Blotting was performed according to standard procedures using primary antibodies as follows: Two Lpd-specific antibodies were purchased from Sigma (HPA016744 recognising the N-terminus, Sigma-Aldrich, Germany) and Santa Cruz Biotechnology (European Support, Heidelberg, Germany; E-12 antibody, epitope unknown). A third Lpd antibody raised against the C-terminus of the protein was as described (Krause et al., 2004). GAPDH antibody (clone #6C5) was purchased from Calbiochem (Merck-Millipore, Germany), beta-actin antibody (ab8227) from Abcam, Germany, and Profilin1 antibody (P7624) from Sigma-Aldrich, Germany. Remaining primary antibodies were as previously published: Polyclonal VASP and PREL1 antisera (Jenzora et al., 2005), and Nap1-B (#4953-B) antiserum (Steffen et al., 2004); WAVE2 (Innocenti et al., 2004) and Abi-1/E3b1 (Biesova et al., 1997) antisera were kindly provided by Dr. Giorgio Scita (IFOM Milan, Italy). Peroxidase-coupled anti-mouse IgG or anti-rabbit IgG secondary antibodies were purchased from Dianova GmbH (Germany).

### Quantification of active Rac proteins

Quantification of GTP-loaded Rac1, 2 and 3 proteins in cell lysates of B16-F1 and Lpd KO cells, grown to 30-50% confluency, was performed using the R*ac1, 2, 3 G-LISA™ Activation Assay* (purchased from Cytoskeleton Inc.), according to manufacturer’s instructions. Results were averaged for 3 independent repetitions, each performed in triplicates for wild-type and Lpd KO cell clones. Functionality of the assay was confirmed with provided positive control (Rac control protein), quantifications of which were excluded from final results.

### Phalloidin stainings and immunolabellings

For phalloidin stainings, cells were fixed for 20 min with a mixture of pre-warmed (37°C) 4% PFA and 0.25% glutaraldehyde, followed by permeabilization in 0.1% Triton-X100/PBS for 1 min. For immunostainings of endogenous proteins, the same procedure was employed except that cells were fixed with 4% PFA alone, and for zyxin stainings, cells were permeabilized first using 0.3% Triton-X100 in 4% PFA/PBS for 1 min followed by fixation for 20 min with 4% PFA/PBS. In case of antibody stainings, samples were blocked with 5% horse serum in 1% BSA/PBS for 30-60 min. ATTO-488- and ATTO-594-labelled phalloidin were purchased from ATTO-TEC GmbH (Germany). The following primary antibodies were used: Mena monoclonal antibody A351F7D9 (Lebrand et al., 2004), monoclonal Abi antibody (clone W8.3), kindly provided by Dr. Giorgio Scita (IFOM Milan, Italy), monoclonal p16A/ArpC5A 323H3 (Olazabal et al., 2002) and monoclonal zyxin antibodies (Rottner et al., 2001). Polyclonal antibodies against murine full-length VASP (residues 1-375) were raised by immunizing a female New Zealand White rabbit with recombinant protein following 5 boosts at two-week intervals by standard procedures.

### Lamellipodial width, actin filament and protein intensity measurements

Lamellipodial F-actin intensity was determined by measuring the average pixel intensities of lamellipodial regions of phalloidin-stained cells, microspikes excluded, with intensities of background regions being subtracted from lamellipodial actin intensity values. Quantitation of Arp2/3 complex intensities (p16A subunit) in the lamellipodia of different cell lines was done as described (Dimchev et al., 2017). For determining lamellipodial intensities of VASP and Abi, intensity scans from 3 pixel-wide lines drawn across lamellipodia at 3 random cellular locations were generated using Metamorph. Upon subtraction of the minimum intensity of each scan defined as background outside the cell, peak intensities from the three individual measurements were averaged and expressed as maximum intensity counts per pixel. Lamellipodial width was quantified with Metamorph using images of phalloidin-stained cells, by drawing lines from lamellipodia tips into more proximal cellular regions up to distal edges of the lamella, followed by measuring their lengths. For each cell, lamellipodial widths in 3 random cellular locations were measured, and values averaged in order to derive the respective single value per cell; obtained values were expressed as μm.

### Fluorescent recovery after photobleaching (FRAP) for investigating protein turnover or actin network polymerization rates

FRAP experiments were performed on an Axio Observer (Carl Zeiss, Jena) using a 100x/1.4NA Plan-Apochromat oil immersion objective. EGFP–VASP and –Abi, localizing at the tip of lamellipodial regions, were bleached by employing the 2D-VisiFRAP Realtime Scanner (Visitron Systems) using 60 mW output power of a 405 nm diode laser (Visitron Systems), in order to achieve nearly complete bleaching for each component at the lamellipodial tip. A time interval of 1500 msec and an exposure time of 500 msec were used for image acquisition. All intensity values were derived by region measurements using Metamorph software. Lamellipodial intensities before and after the bleach were corrected for background and acquisition photobleaching using regions outside of and inside the cell, respectively, and processed using Microsoft Excel. Data were fitted in SigmaPlot 12.0 (Scientific Solutions SA, Pully-Lausanne, Switzerland) using dynamic curve fits for exponential rise to maximum, and half-times of recovery calculated using equation t_1/2_=-1/b*ln(0.5), as described (Dimchev and Rottner, 2018; Steffen et al., 2013).

Lamellipodial actin polymerisation rates in B16-F1 wildtype and Lpd knockout cells were obtained by photobleaching incorporated, EGFP-tagged actin. A region exceeding the lamellipodial area was bleached in each case. This allowed monitoring the recovery of fluorescence in the entire actin network over time, essentially as described previously (Dimchev and Rottner, 2018; Kage et al., 2017; Steffen et al., 2013). Final actin network assembly rates were expressed as µm/min.

### Cell spreading quantification of adhesion area

Cells were seeded onto coverslips pre-coated with either 25 μg/ml laminin or 25 μg/ml fibronectin (as described above). 15 min, 60 min or 3 hrs following cell seeding, coverslips were fixed with 4% PFA / 0.25% glutaraldehyde, permeabilized with 0.1% Triton-X100/PBS for 1 min and stained for F-actin with phalloidin (as described above). Images were taken with a 40x/1.3NA Apochromat oil objective. Cell area measurements were derived from drawing cell contours encompassing the full cell area with ImageJ, following calibration to respective objective magnification. Final data were averaged for all cells measured and displayed in µm^2^. Quantification of adhesion area in cells stained for zyxin was performed using ImageJ by manually adjusting appropriate thresholds and employing the particle analysis plugin. For enhancing accuracy, the area around the nuclear region, characterised with nonspecific background fluorescence, was excluded from measurements. Since it was challenging to separate individual adhesions in computer-aided analyses, we presented data as average, adhesion-occupied areas in percent of whole cell area.

### Nascent adhesion quantification

Cells were transfected with EGFP-paxillin or co-transfected with EGFP-paxillin and pRK5-myc-Rac1 (wt) in case of Fig. S5B, and dual-channel time lapse movies (EGFP fluorescence and phase contrast) acquired on an Axio observer microscope (Carl Zeiss, see above) with 100x/1.3NA Neofluar oil immersion objective and 20 sec time interval between frames. Time-lapse movies were opened in Metamorph and brightness and contrast adjusted to optimize visualisation of lamellipodia and nascent adhesions. Lamellipodia width could be determined in both fluorescence and phase-contrast channels, due to these structures appearing as slightly brighter and darker in respective channels, which was due to the increased concentration of actin filaments in this region. Contours encompassing the entire width of lamellipodial regions were manually drawn on Metamorph for each time frame measured, and further separated into back and front lamellipodial regions. This was implemented by drawing multiple perpendicular lines connecting front and back of the lamellipodial region at multiple locations, and marking the middle of each perpendicular line. By connecting the middle parts of all perpendicular lines with the Metamorph multi-line tool then allowed separating the lamellipodium into two essentially equal halves. Nascent adhesions in lamellipodia were manually counted in 4 consecutive frames, and individually for back and front parts of the lamellipodium. For each cell, the number of adhesions in each region measured (either front or back) were averaged for the 4 consecutive frames, thus deriving the average number of adhesions present in each region over the course of 80 seconds. Data were presented as number of nascent adhesions per 10 µm^2^, grouped into lamellipodial phenotype categories defined above (smooth, intermediate and chaotic), and displayed separately for back and front parts of lamellipodia.

### Vaccinia virus infections

Cell lines were infected with the WR strain of vaccinia virus and processed for immunofluorescence as previously described (Arakawa et al., 2007a; Arakawa et al., 2007b). In case of HeLa, cells were fixed at 9 hr post-infection and for KO MEFs at 15 hr post-infection. The following antibodies were used for immunofluorescence: Nck (Millipore), N-WASP (Moreau et al., 2000), VASP (BD Transduction Laboratories), Lpd (Krause et al., 2004), B5 (Hiller and Weber, 1985). Actin tails were stained with Texas Red or Alexa-488 phalloidin (Invitrogen), and viral DNA was observed using DAPI staining. The quantification of number of actin tails and number of extracellular virus was performed through manual counting, all other analyses were performed as described previously (Humphries et al., 2014). For all quantifications, counts were performed over multiple cells in three separate experiments, with the n in each case given in the figure legend. Graphs and statistics were compiled using Prism (Graphpad software, USA), comparison of two data sets was carried out using an unpaired t-test.

### Cell edge tracking for lamellipodial morphodynamic analysis

Cells were transfected with EGFP-CAAX construct or EGFP-Rac1 (wt) in case of the data shown of Fig. S5C-E, and time-lapse movies acquired on an inverted microscope (Axiovert 100 TV; Carl Zeiss) using a 100x/1.4NA Plan-apochromatic oil immersion objective, with a time interval of 1 second. To enhance cell edge detection, acquired fluorescent time-lapse movies were processed on ImageJ with *Smooth, Gaussian blur*, *Enhance contrast* and *Find edges* filters. Brightness and contrast levels were further adjusted to obtain optimal separation of edge from background. Cell edge analysis was initiated with ImageJ by using the *JFilament* 2D plugin (Smith et al., 2010b). The following *JFilament* parameters were found optimal for B16-F1 cell edge detection with aforementioned acquisition settings: Alpha=15, Beta=10, Gamma=4000, Weight=0.3, Stretch Force=15, Deform iterations=400, Point Spacing=5, Image Smoothing=1.01. 120 frames (i.e. 2 min of acquisition time) were processed in order to obtain “snake” files (or snakes), each one containing the spatiotemporal positions of the tracked cell contour, defined by x and y coordinates of a given number of points. Snake files were processed in Matlab R2017a (Mathworks, Massachusetts, United States), in order to derive velocity, curvature maps and specific morphodynamic parameters.

### Matlab script and displacement field of the contour

Following cell edge tracking and obtaining contour coordinates from Jfilament for every frame, a Matlab script was employed to calculate the displacement and velocity fields, using a method aiming to determine the correspondences of points on the measured cell edge contour over subsequent frames. We employ uniform springs and normal springs to find the corresponding positions of points between the curve in time step *t* (i.e. *C_t_*), and the curve in time step *t+1* (i.e. *C_t+1_*) where *t* is an arbitrary time frame. Uniform springs connect neighboring points of a contour C_t+1_. Normal springs connect these points to the points representing the continuation of the normal direction of the contour C_t._ For regions where curves in two subsequent frames are parallel, the assumption is that each point is moved in the normal direction and the correspondence between curves is achieved by following the normal vectors from C_t_, to C_t+1_. For regions of high deformations and enhanced convexity or concavity in one of the curves, meaning the curves not being parallel to each other, a normal mapping would not be reliable. In these situations, uniform mapping is employed resulting in evenly spaced distribution of points on C_t+1_. In order to add an adaptivity feature to the correspondence algorithm, normal and uniform mappings along the curves were combined, but with variable weights, the latter being defined as a function of local deformation or parallelism degree. The variability of weights enabled constructing the basis of the mapping between two curves, relying on the normal mapping method in the regions of low deformation, followed by using the uniform mapping to interpolate the mapping in the regions of high deformation. In order to implement this principle, a mechanical spring model was employed similar to the one described previously (Machacek and Danuser, 2006), containing two types of springs: springs of normal or uniform mapping. Unlike the mechanical model used by Machacek and Danuser (2006), in our model the springs responsible for normal mapping were also considered as linear springs and all the springs are assumed to be bound to the cell boundary. This simplifies the nonlinear spring system to a linear one in the 1D arc length coordinate, and facilitates the computations. At each time point t, there are N fixed nodes Q_i_, i= 1:N on C_t_, whose coordinates are obtained by applying the same procedure in the previous time frame, and the aim of the algorithm is to find the position of their corresponding nodes P_i_, i= 1:N on C_t+1_.

Nodes on C_t+1_ are connected by two kinds of springs. Uniform mapping springs connect neighbouring nodes and their rest length is defined as l_0 t+1_=L_t+1_/N, where L_t+1_ is total length of the curve C_t+1_.

Normal mapping springs connect each node P_i_ to its normally mapped position M_i_, i= 1:N, which is defined as the intersection of curve C_t+1_ and normal vector from Q_i_ on curve C_t_ (see figure). The rest length of these springs is zero, thus they tend to attract nodes toward their normally mapped positions M_i_. The strength of this attraction depends on the spring constant, the latter depending on deformation degree.

Degree of deformation at each node P_i_ was defined as:

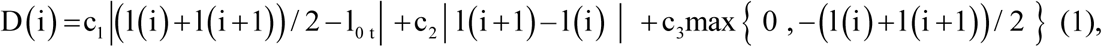

where c_1_, c_2_, c_3_ are constants and l(i) is the length of the segment of C_t+1_ between M_i-1_ and M_i_, which can be negative if M_i_ appears before M_i-1_.

The first term in aforementioned equation decreases in the regions where curves C_t_ and C_t+1_ are close to parallelism, i.e. it measures the similarity of the direction of curves C_t_ and C_t+1_ at any point i. The second term captures the similarity of the curvature of curves C_t_ and C_t+1_ at any point i. The third term penalises the regions where the order of M points does not match the order of Q nodes. In these regions, the additional increase in deformation degree caused by the third term reduces the possibility of topological violations, which is the difference between the order of P nodes and Q nodes.

Spring constants for normal mapping springs and uniform mapping springs were defined as functions of the degree of deformation:

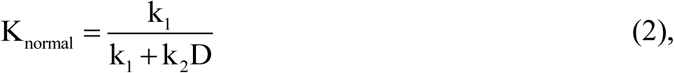

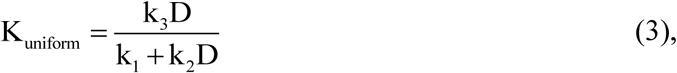

where k_1_, k_2_ and k_3_ are constants.

These relations indicate the predominance of uniform and normal mapping in, respectively, high and low deformation regions.

For each time point t, a system of N nodes on C_t+1_ was constructed, which are connected by different springs, as explained above. In order to find unknown positions of nodes P_i_, N force balance equations were solved. By using the positions of nodes Q_i_ and their corresponding nodes P_i_, the displacement and velocity fields of the curve C_t_ were found. The effects of curve ends on the mapping were taken into consideration. The result of the mapping, especially in the regions close to ends, can depend on the position of end points of the curves in different time frames, and in order to compensate for this dependency, 5% of the snake lengths at both ends of tracked contours were removed. The Matlab Script will be available upon request.

### Quantitative analysis of cell edge morphodynamic metrics and PCA analysis

Finding the corresponding location of each point on the tracked cell edge allowed deriving the trajectory of each point over time and obtaining the velocity of each point between different frames. Once the local velocities of points along the cell boundary were calculated, velocity maps were constructed by plotting velocity data in a 2D coordinate system, with time and position along the cell boundary defined as x- and y-axis, respectively. The velocity at any point on curve and any time was indicated by colours, as shown in Figure 3. For the morphodynamic behaviour of the leading edge, only the normal component of the velocity was considered. Similarly, maps for curvature along the cell boundary in time were constructed. Since the curve in our model is a piecewise linear curve represented by a finite number of points, the local curvature at any point was calculated using the position data of the point and its two neighbouring points. The velocity and curvature maps contain the majority of morphodynamical information of the analysed edge. The morphodynamical behaviour of cells was subsequently quantified by defining descriptors based on these maps (explained in Supplementary Table S1). Principal Component Analysis (PCA) was performed on the defined morphodynamical metrics, using Matlab.

### Image Processing and Statistical analysis

Brightness and contrast levels were adjusted using MetaMorph software v7.7.8.0. Figures were processed and assembled on Adobe Photoshop CS4 (Adobe Systems, San Jose, CA). Data analyses were performed with MetaMorph, Fiji or ImageJ, Microsoft Excel 2010 and Sigma plot 12.0 (Systat Software, Erkrath, Germany). Data sets were routinely repeated three or more independent times, and statistically compared with paired t-tests (if normally distributed) or alternatively, non-parametric Mann–Whitney rank sum tests using Sigma plot. Probability of error of 5% or less (\**p* < 0.05) was considered as statistically significant.

## ACKNOWLEDGMENTS

We would like to thank Drs. Giorgio Scita, Laura Machesky and Wolfgang Kranewitter for reagents, and Dr. Naoko Kogata (The Francis Crick Institute, London, UK) for help with assembling Vaccinia-related figures, Dr. Frieda Kage (Geisel School of Medicine at Dartmouth, Hanover, NH, USA) for fruitful discussions, and Brigitte Denker for excellent technical assistance.

## Supplementary information

**Fig. S1.**
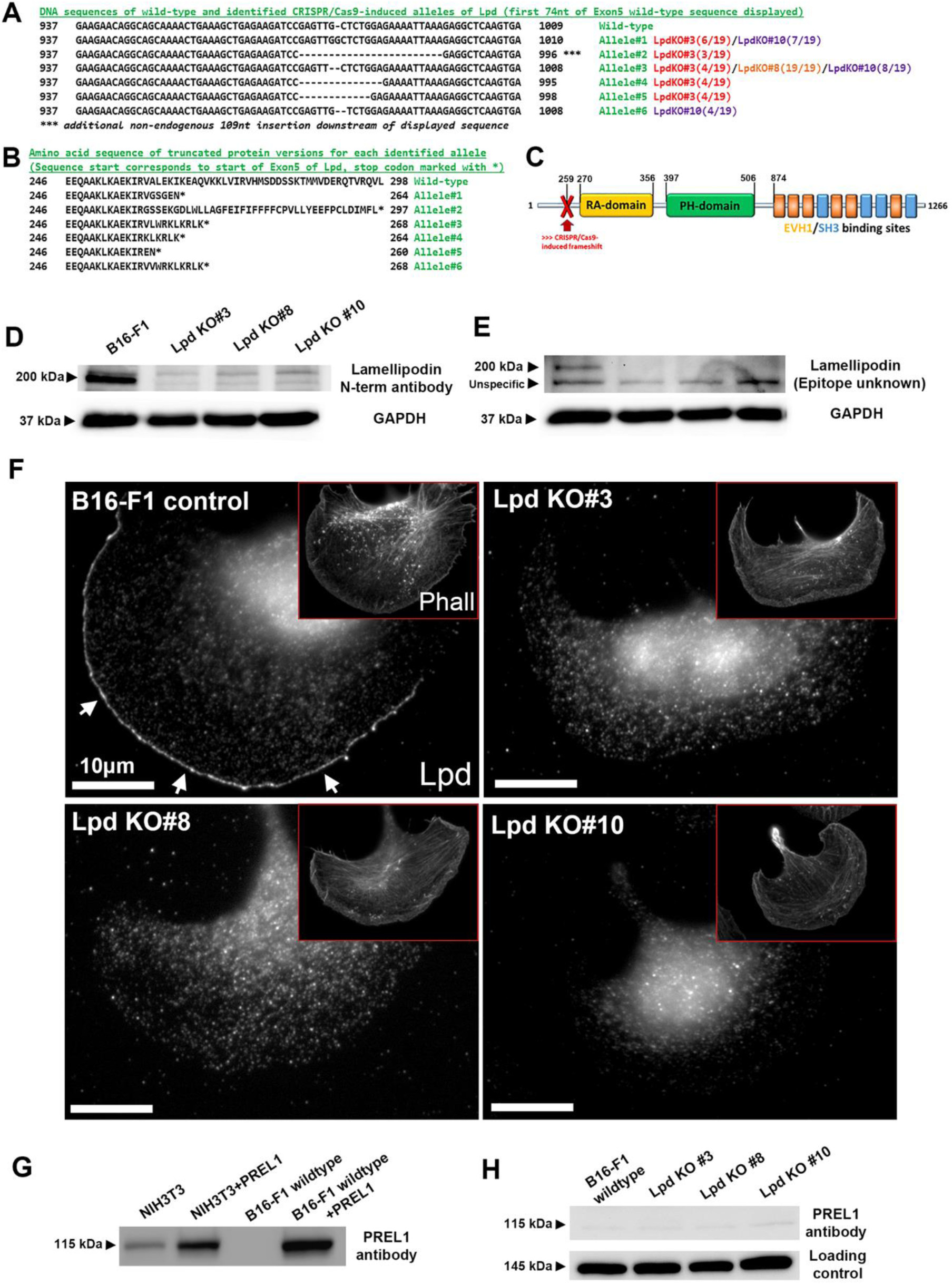
Confirmation of lack of Lpd expression in Lpd KO lines as well as of RIAM/PREL1. (A) DNA sequence alignments of wildtype Lpd Exon-5 fragment *versus* all alleles discovered in B16-F1 Lpd KO cell clones by DNA sequencing, including respective mutations. Different colours indicate either Lpd KO#3 (red), Lpd KO#8 (orange) or Lpd KO#10 (purple). Numbers in brackets following each cell clone indicate the frequency of occurrence of respective allele. (B) Protein sequence alignments of wildtype Lpd Exon-5 fragment *versus* all alleles discovered in Lpd KO cell clones. Allele numbers correspond to those in A. * Indicate the occurrence of a CRISPR/Cas9-induced premature stop codon. (C) A graphical representation of the Lpd domain structure, with CRISPR/Cas9 target site labelled in red (not drawn to scale), indicating that truncated versions of the protein (shown in B) would lack every domain located downstream and reportedly essential for its physiological function and interaction with other proteins. (D, E) Immunoblots with extracts from wildtype and Lpd KO cell lines using two alternative Lpd antibodies, as indicated; GAPDH was used as loading control. Note the absence of detectable Lpd protein levels in all characterized, independently generated B16-F1 knockout clones (Lpd KO#3, -#8 and -#10). (F) Immunolabellings using Lpd antibodies and counterstaining with phalloidin (insets) in distinct, indicated B16-F1 cell lines. White arrows point at clear accumulation of Lpd at the front edge of the lamellipodium in B16-F1 control cells, as expected, a staining pattern completely absent in representative cells shown for each KO cell population, in spite of the presence of a clear lamellipodium in each case (see phalloidin insets, Phall). (G) Immunoblot with RIAM/PREL1-specific antibody using lysates from NIH3T3 fibroblasts or B16-F1 cells, either untransfected or transfected with an untagged version of full-length PREL1. No detectable protein expression was observed in B16-F1 wildtype cells, as opposed to NIH3T3 or respective cells overexpressing PREL1 protein. Ponceau red staining was used as loading control (not shown). (H) Immunoblotting of B16-F1 wildtype and derived Lpd KO clones to test for potential upregulation of expression of RIAM/PREL-1 upon Lpd removal. However, at best a very faint band at expected molecular weight (115kDa) was detectable in all cell lines tested. A non-specific cross-reaction of the antibody was used as loading control.

**Fig. S2.**
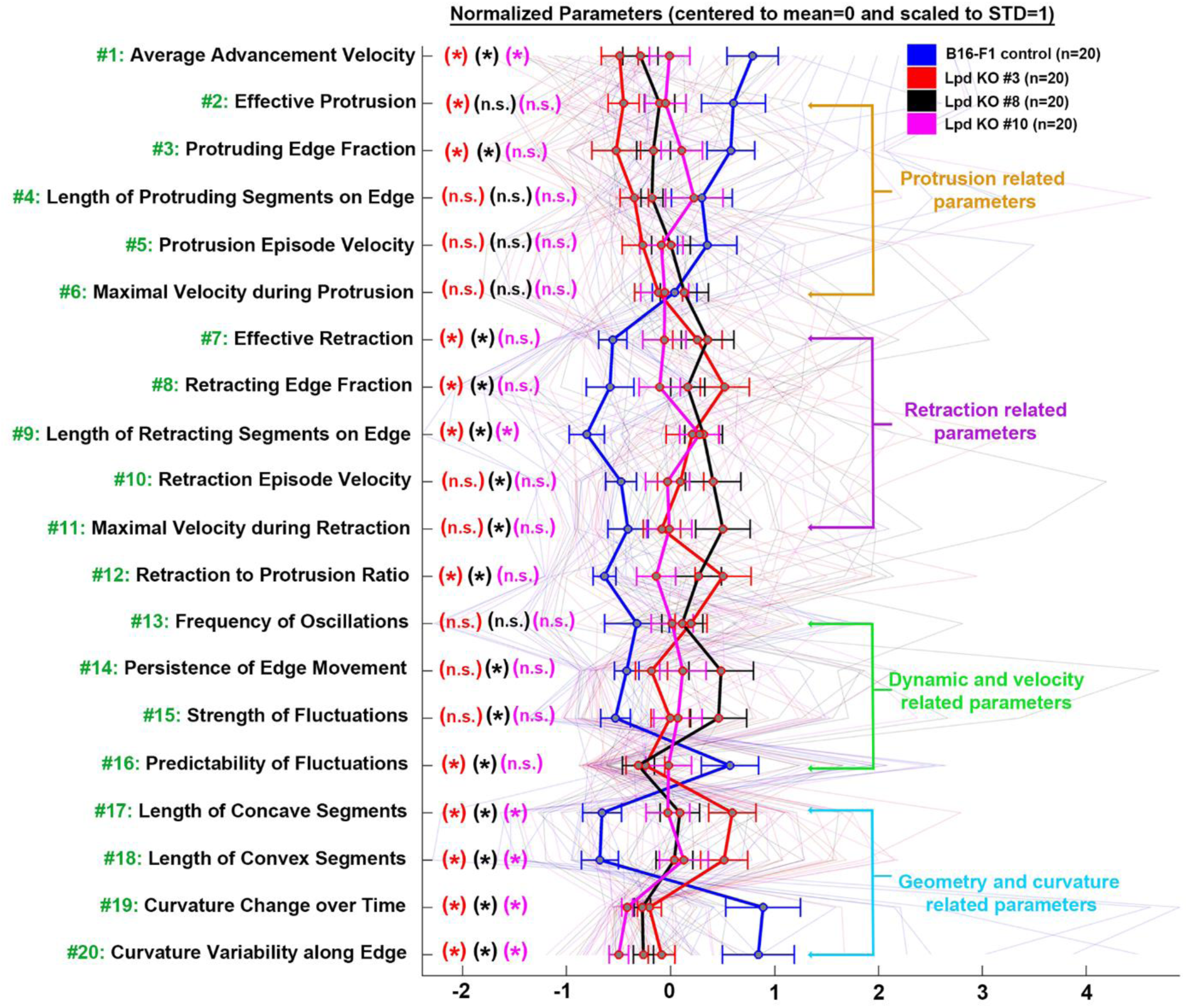
Morphodynamic protrusion analysis with B16-F1 wildtype and characterized Lpd knockout clones displayed individually. Data displayed as described for Fig. 4, except that knockout clones are displayed separately.

**Fig. S3.**
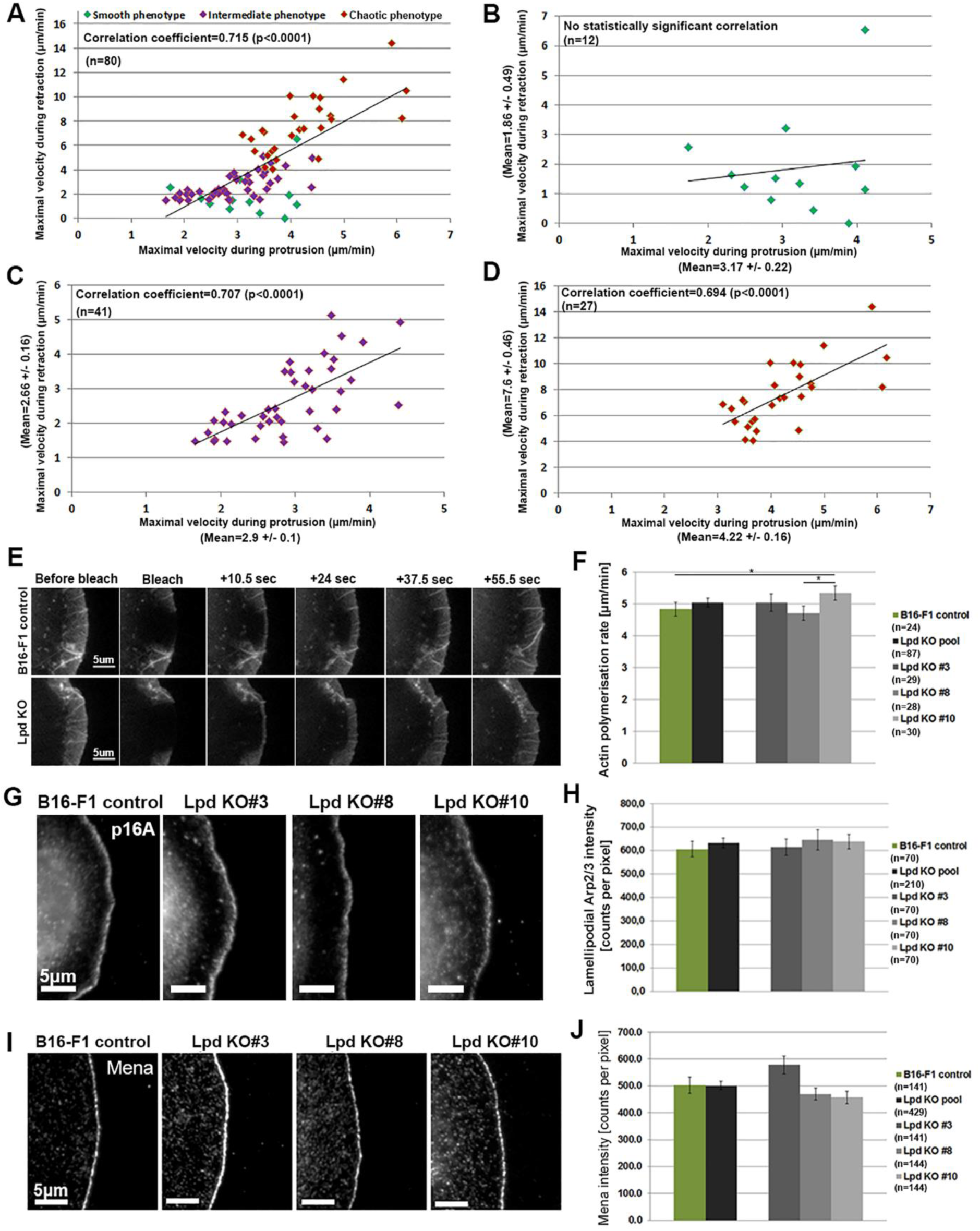
Protrusion/retraction correlation analysis and assessment of lamellipodial actin assembly or Arp2/3 and Mena intensities. (A) B16-F1 cells grouped into the three lamellipodial protrusion categories defined in Fig. 3 were plotted into a 2D-coordinate system with values for maximal velocity during protrusion and maximal velocity during retraction displayed on x- and y-axis, respectively. B16-F1 wildtype and Lpd KO cells were pooled for this analysis. The data reveal a statistically significant, positive correlation between the two parameters. (B-D) Correlation analyses as described in A, but with protrusion categories displayed separately, as indicated. Note the absence of a statistically significant correlation in the smooth protrusion phenotype, caused perhaps by the great variability and/or low frequency of retraction events in this category. All data are arithmetic means ± SEM. n equals number of individual movies and cells analysed. (E, F) Rates of actin network polymerization in the lamellipodium as assessed by FRAP of ectopically expressed, EGFP-tagged beta-actin in B16-F1 control *versus* Lpd knockout cells, as indicated (F). Quantitation of Lpd KO results is either displayed individually or as pooled population (n equals number of cells analysed). One out of three Lpd KO clones (#10) appeared to display even a slight, but statistically significant increase in lamellipodial actin assembly rate (* indicates p<0.05), although this clonal variability was eliminated in the KO pool population. (G) Immunostaining and (H) quantitation of Arp2/3 complex (subunit p16A/ArpC5A) in B16-F1 control and Lpd knockout cells, as indicated. n=number of cells analysed, and data are arithmetic means ± SEM. Statistics revealed no significant differences between any pair of experimental groups (not shown). (I) Immunostaining and (J) quantitation of Mena intensity at the lamellipodium edge in B16-F1 control and Lpd KO cells, as indicated. In spite of moderate clonal variability among individual Lpd KO clones, no statistically significant differences between genotypes were found (not shown).

**Fig. S4.**
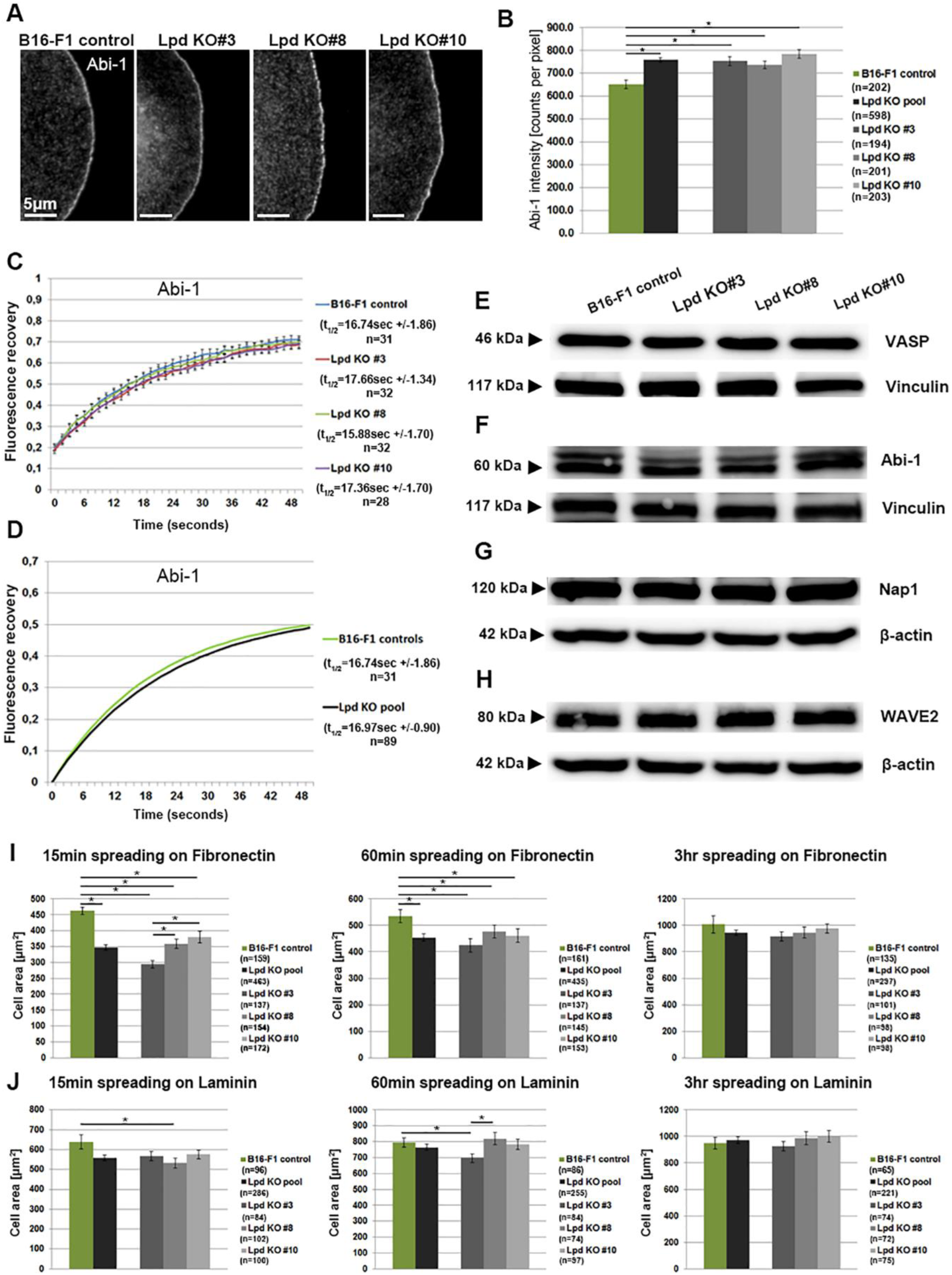
Lpd-KO increases lamellipodial Abi-1 accumulation and reduces cell spreading. (A) Representative images of immunolabelling with Abi antibody at lamellipodia edges of B16-F1 control and Lpd knockout cells, as indicated. (B) Quantitation of lamellipodial Abi intensities in B16-F1 wildtype *versus* Lpd knockout cells. Data are arithmetic means ± SEM, asterisks above bar charts indicate statistically significant differences between experimental groups, p<0.05, n equals number of cells measured. (C) Averaged raw data of fluorescent recovery curves after photobleaching of EGFP-tagged Abi-1, plotted for B16-F1 cells and all three individual Lpd KO clones. Determined half-times of recovery (t_1/2_, in seconds, as derived from fitted data, see D and not shown) for each group are displayed on the right, n equals number of FRAP movies used. (D) Curve fits of averaged, fluorescent recovery data of photobleached, lamellipodial Abi expressed in B16-F1 wildtype *versus* Lpd KO cells (all lines pooled for the latter). Derived half-times of recovery are displayed on the right, confirming the absence of any significant difference in Abi turnover rates between distinct cell types. (E-H) Western blots with total protein lysates using antibodies specific for VASP (E) or the WRC subunits Abi-1 (F), Nap1 (G), and WAVE2 (H). Equal loadings were controlled in each case by Ponceau S staining (not shown), and antibodies specific for vinculin or β-actin, as indicated. (I, J) B16-F1 wildtype cells or Lpd KO clones were seeded onto fibronectin (I) or laminin (J) for time periods of 15, 60 or 180 min, as indicated, fixed and quantified for cell area covered. Note the statistically significant reduction in cell area in the Lpd KO cell pool as compared to controls observed upon 15 or 60 min of spreading on fibronectin (I). The same trend was also observed after seeding on laminin, albeit less pronounced and not in a statistically significant fashion for the Lpd KO cell pool (J). Irrespective of substratum, this phenotype appeared to be compensated for upon extended spreading periods (180 min). Data are arithmetic means ± SEM, and single asterisks above bar charts show statistically significant differences between designated groups (* indicates p<0.05); n equals numbers of cells measured.

**Fig. S5.**
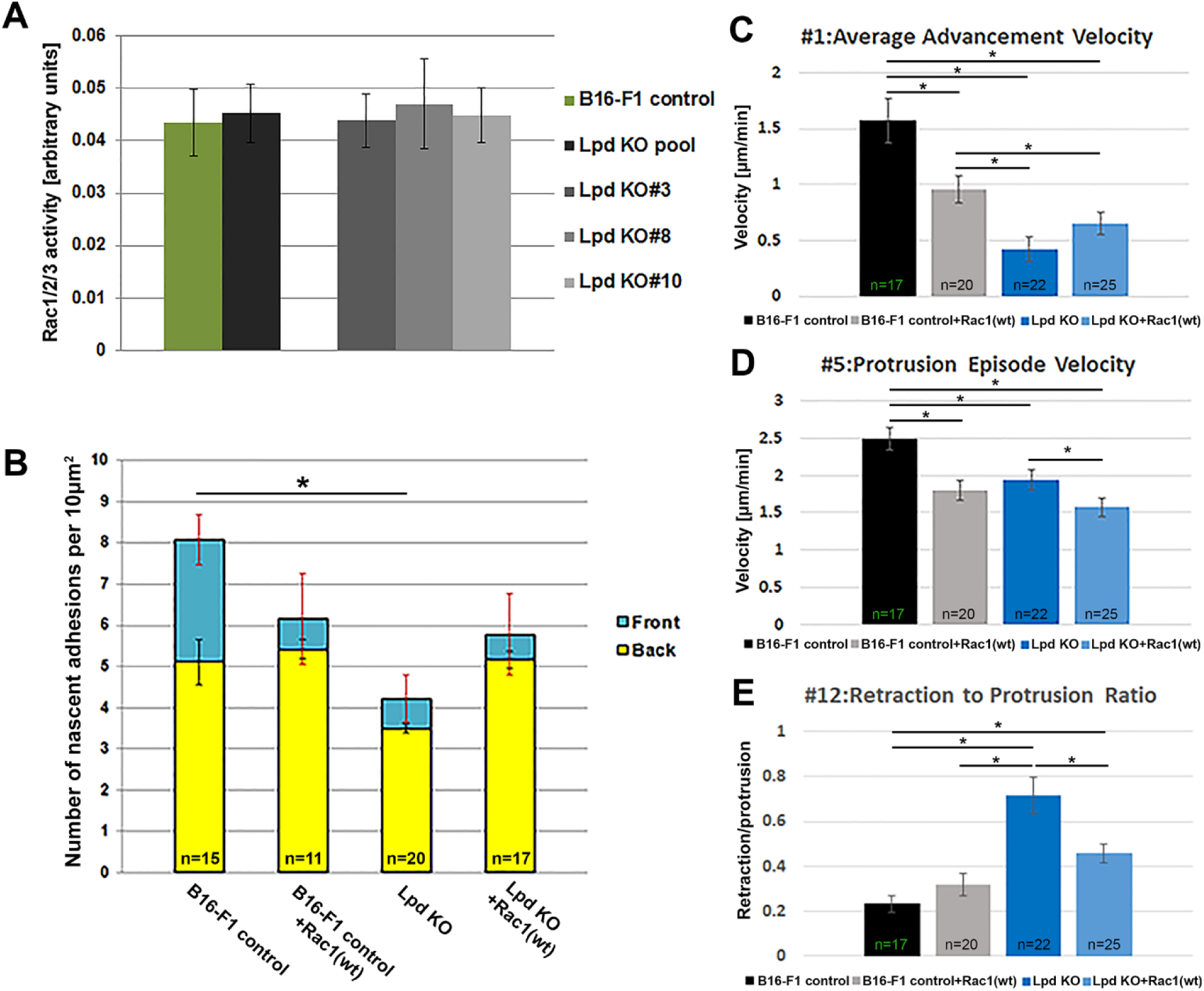
Overexpression of Rac1 (wt) protein modulates nascent adhesion formation and cell edge morphodynamic behaviour. (A) Amount of active (GTP-loaded) Rac1, 2 and 3 proteins do not differ significantly between B16-F1 wildtype and Lpd KO cell clones, as quantified by G-Lisa colorimetric assay. Mean values for each condition are calculated from 3 independent experiments, each consisting of 3 replicates. (B) Quantification of numbers of nascent, EGFP-paxillin-containing adhesions in lamellipodia of B16-F1 wildtype or Lpd KO#3 cells, with or without ectopic Myc-Rac1(wt) expression. Quantifications are also separated for front and back lamellipodial regions, highlighted as cyan and yellow bars, respectively. (C, D, E) Bar plots of selected morphodynamic parameters comparing their mean values between B16-F1 wildtype and Lpd KO#3 cells, with or without ectopic expression of EGFP-tagged Rac1(wt). The plasma membrane localization of EGFP-Rac1 was simultaneously employed as fluorescence marker enabling morphodynamic analysis, whereas controls expressed inert EGFP-CAAX (also see Methods). Parameter numbers above each plot correspond to those in Supplementary Table 1 and Figs. 4, S2 and S6. Data are arithmetic means ± SEM, and single asterisks above bar charts show statistically significant differences between designated groups (* indicates p<0.05); n equals numbers of cells measured.

**Fig. S6.**
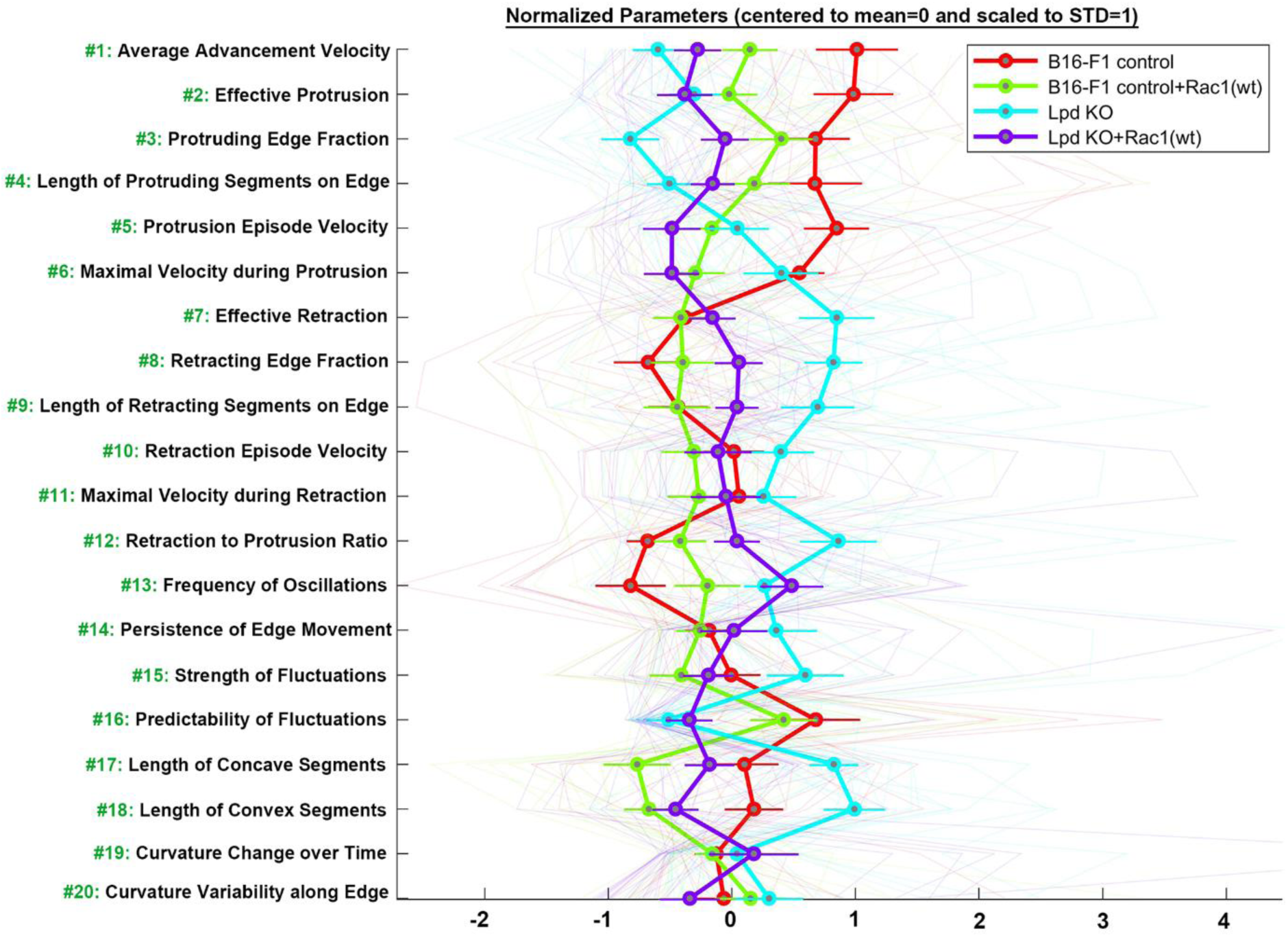
Results from morphodynamic protrusion analysis of wildtype *versus* Lpd knockout cells with or without ectopic Rac1-expression. Data displayed as described for Fig. 4 and Fig. S2, except that B16-F1 wildtype or Lpd KO clone #3 were transfected with EGFP-Rac(wt), as indicated, or with EGFP-CAAX as control.

**Fig. S7.**
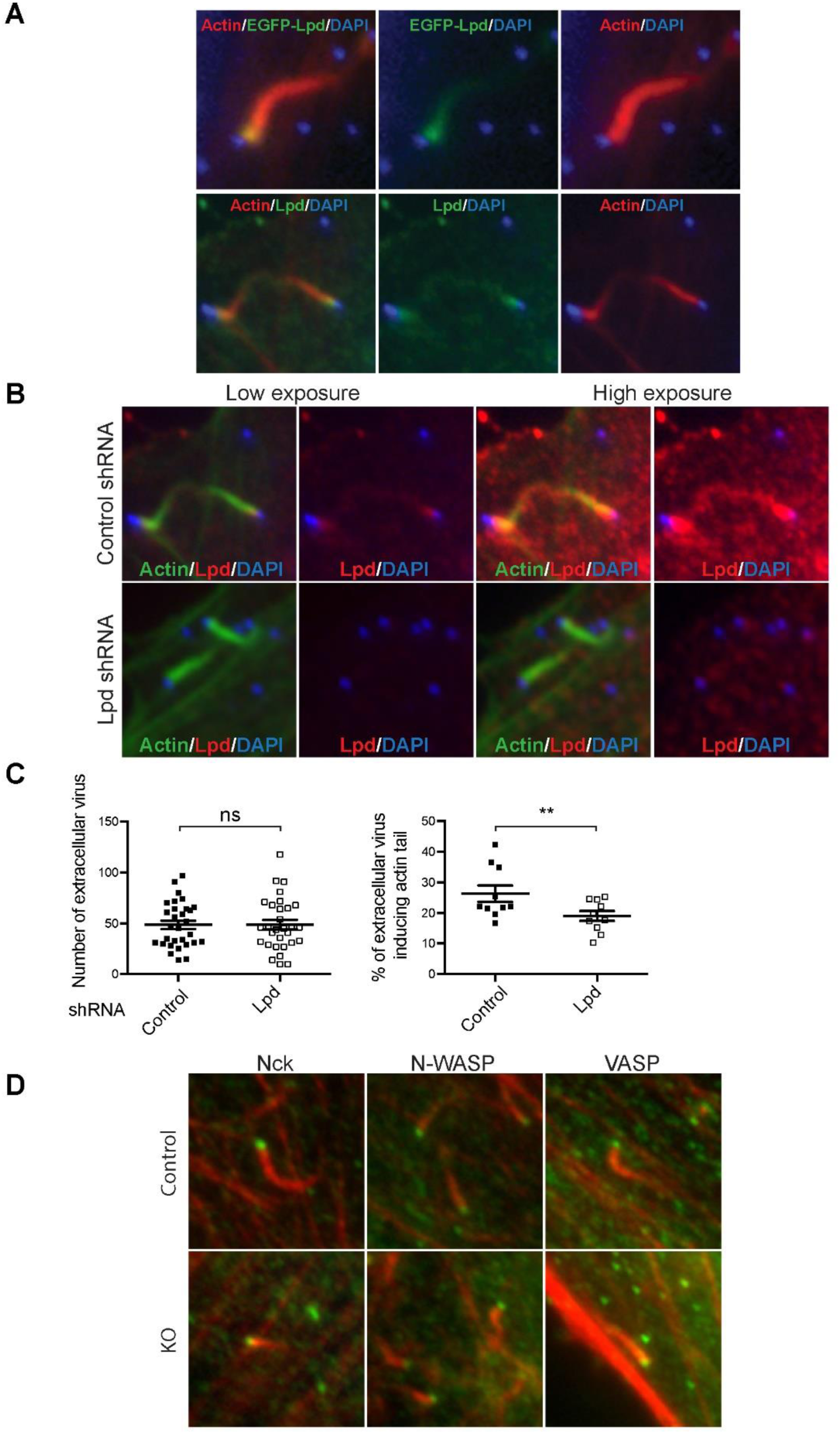
Lpd localisation below Vaccinia virus and lack of effects on extracellular virus positioning and actin machinery recruitment. (A) Immunofluorescence images showing that both EGFP-tagged and endogenous Lpd (green) are recruited to the tip of actin tails (red) induced by Vaccinia virus (blue) in infected HeLa cells. (B) Immunofluorescence images reveal that after shRNA treatment of Hela cells, small amounts of endogenous Lpd (red) can still be observed at the tips of actin tails (green). Note this localisation can only be observed upon significant enhancement of fluorescence intensity post imaging (“high exposure”). (C) Quantification of number of extracellular virus per cell (n=30), and percentage of extracellular virus inducing an actin tail (n=30), in control and Lpd shRNA-treated cells. (D) Immunofluorescence images reveal the presence of endogenous core actin assembly factors on Vaccinia surfaces, Nck, N-WASP as well as the Lpd-ligand VASP (green) at the tips of vaccinia-induced actin tails (red, phalloidin), in control (Lpd fl/fl) and Lpd knockout MEFs (KO).

**Fig. S8.**
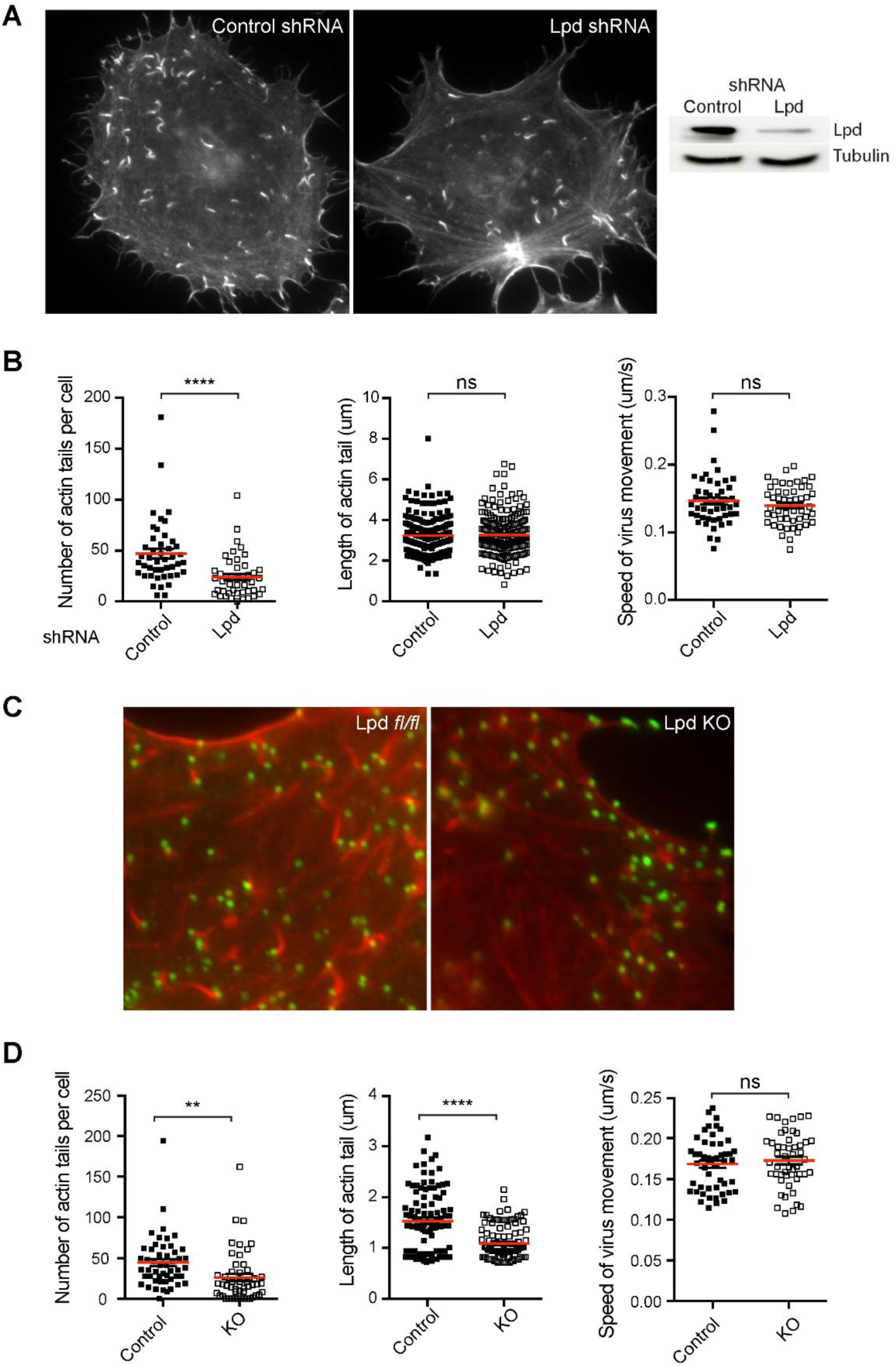
Frequency of Vaccinia actin tail formation, but not their rate of motility is reduced by Lpd depletion. (A) Representative fluorescence images of HeLa cells treated with control and Lpd shRNA. In both cases, phalloidin staining reveals actin tails induced by vaccinia virus. The reduction in Lpd levels by shRNA was confirmed by immunoblot. (B) Quantification of the number of actin tails per cell (n=45 cells), the length of actin tails (n=200 tails), and the speed of viral movement (n=50 viruses), in control and Lpd shRNA-treated cells. (C) Fluorescence images of actin tails (red, phalloidin) produced by vaccinia (green, DAPI) in parental and Lpd KO MEFs, as indicated. (D) Quantification of the number of actin tails per cell (n=50 cells), the length of actin tails (n=100 tails), and the speed of viral movement (n=50 viruses), in control (Lpd fl/fl) and Lpd KO MEFs. Red lines correspond to arithmetic means, and error bars SEM from three independent experiments. ****P < 0.0001, **P < 0.01, n.s., not significant.

## Supplementary Table

**Table S1.**
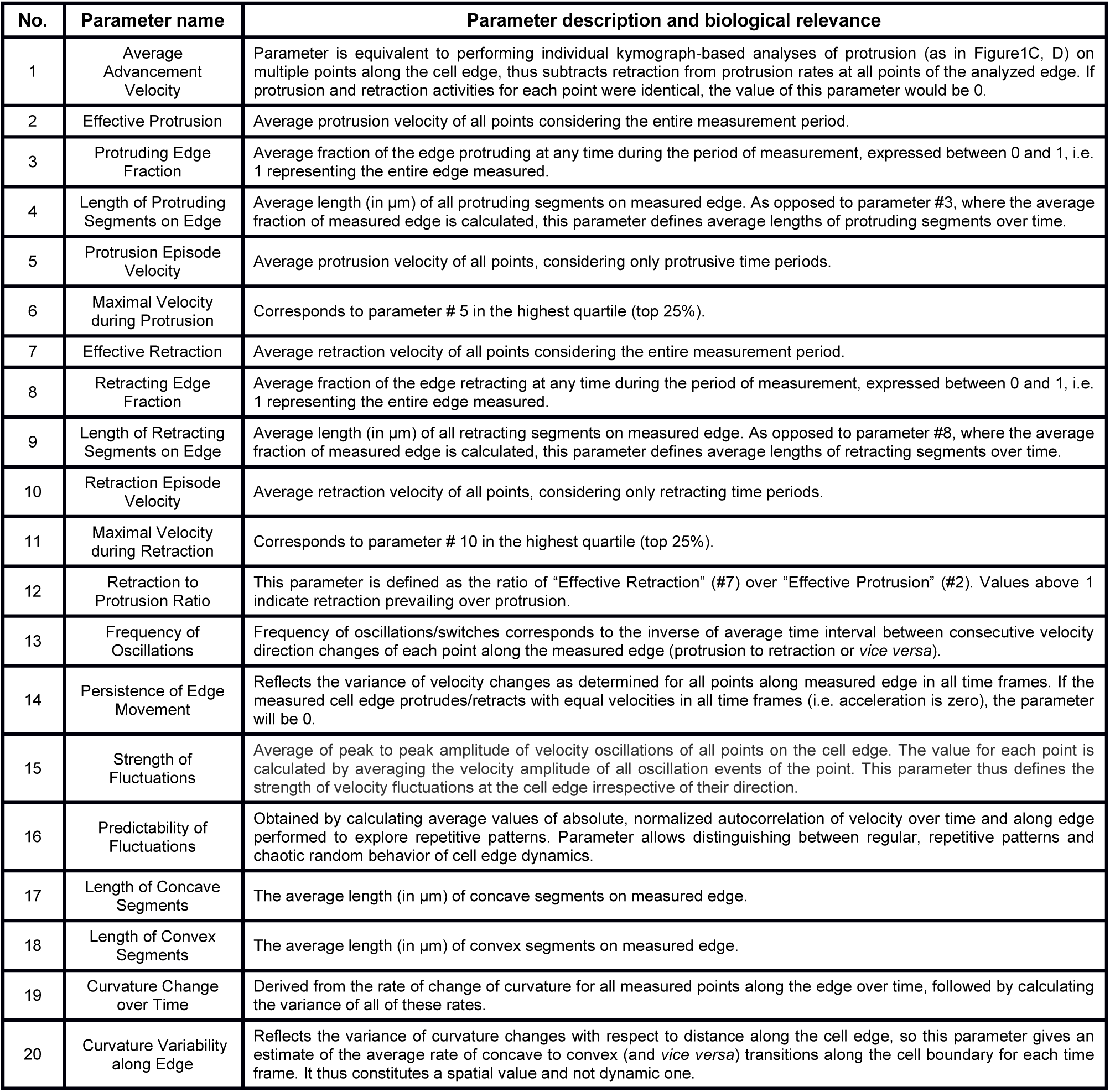

| No. | Parameter name | Parameter description and biological relevance |
| --- | --- | --- |
| 1 | Average Advancement Velocity | Parameter is equivalent to performing individual kymograph-based analyses of protrusion (as in Figure1C, D) on multiple points along the cell edge, thus subtracts retraction from protrusion rates at all points of the analyzed edge. If protrusion and retraction activities for each point were identical, the value of this parameter would be 0. |
| 2 | Effective Protrusion | Average protrusion velocity of all points considering the entire measurement period. |
| 3 | Protruding Edge Fraction | Average fraction of the edge protruding at any time during the period of measurement, expressed between 0 and 1, i.e. 1 representing the entire edge measured. |
| 4 | Length of Protruding Segments on Edge | Average length (in $\mu\text{m}$) of all protruding segments on measured edge. As opposed to parameter #3, where the average fraction of measured edge is calculated, this parameter defines average lengths of protruding segments over time. |
| 5 | Protrusion Episode Velocity | Average protrusion velocity of all points, considering only protrusive time periods. |
| 6 | Maximal Velocity during Protrusion | Corresponds to parameter # 5 in the highest quartile (top 25%). |
| 7 | Effective Retraction | Average retraction velocity of all points considering the entire measurement period. |
| 8 | Retracting Edge Fraction | Average fraction of the edge retracting at any time during the period of measurement, expressed between 0 and 1, i.e. 1 representing the entire edge measured. |
| 9 | Length of Retracting Segments on Edge | Average length (in $\mu\text{m}$) of all retracting segments on measured edge. As opposed to parameter #8, where the average fraction of measured edge is calculated, this parameter defines average lengths of retracting segments over time. |
| 10 | Retraction Episode Velocity | Average retraction velocity of all points, considering only retracting time periods. |
| 11 | Maximal Velocity during Retraction | Corresponds to parameter # 10 in the highest quartile (top 25%). |
| 12 | Retraction to Protrusion Ratio | This parameter is defined as the ratio of "Effective Retraction" (#7) over "Effective Protrusion" (#2). Values above 1 indicate retraction prevailing over protrusion. |
| 13 | Frequency of Oscillations | Frequency of oscillations/switches corresponds to the inverse of average time interval between consecutive velocity direction changes of each point along the measured edge (protrusion to retraction or *vice versa*). |
| 14 | Persistence of Edge Movement | Reflects the variance of velocity changes as determined for all points along measured edge in all time frames. If the measured cell edge protrudes/retracts with equal velocities in all time frames (i.e. acceleration is zero), the parameter will be 0. |
| 15 | Strength of Fluctuations | Average of peak to peak amplitude of velocity oscillations of all points on the cell edge. The value for each point is calculated by averaging the velocity amplitude of all oscillation events of the point. This parameter thus defines the strength of velocity fluctuations at the cell edge irrespective of their direction. |
| 16 | Predictability of Fluctuations | Obtained by calculating average values of absolute, normalized autocorrelation of velocity over time and along edge performed to explore repetitive patterns. Parameter allows distinguishing between regular, repetitive patterns and chaotic random behavior of cell edge dynamics. |
| 17 | Length of Concave Segments | The average length (in $\mu\text{m}$) of concave segments on measured edge. |
| 18 | Length of Convex Segments | The average length (in $\mu\text{m}$) of convex segments on measured edge. |
| 19 | Curvature Change over Time | Derived from the rate of change of curvature for all measured points along the edge over time, followed by calculating the variance of all of these rates. |
| 20 | Curvature Variability along Edge | Reflects the variance of curvature changes with respect to distance along the cell edge, so this parameter gives an estimate of the average rate of concave to convex (and *vice versa*) transitions along the cell boundary for each time frame. It thus constitutes a spatial value and not dynamic one. |

## Supplementary Movie Legends

**Supplementary Movie 1. Comparison of protrusion dynamics in control and Lpd-deficient B16-F1 cells.** Phase-contrast movies of B16-F1 wildtype and Lpd knockout cells (Lpd KO#10) migrating on laminin. Note individual examples of Lpd knockout cells, the protrusion of which is characterized by a fluctuating, chaotic mode of lamellipodial dynamics (white arrows).

**Supplementary Movie 2. Exemplary generation of a velocity map.** A simulated contour (yellow line) consisting of multiple points is slowly protruding to the right (black arrow on top) while fractions of the edge are retracting and protruding with differential timing and velocity (see colour-velocity bar on the left). The velocity map on the right is building up in temporal synchrony with contour movements on the left, illustrating how cell edge movements correlate with the outcome of the velocity map.

**Supplementary Movie 3. Exemplary generation of a curvature map.** A map reflecting the spatiotemporal changes of curvature of a simulated contour (from convex to linear and concave, as shown on the colour bar on the left) is generated. Everything is as described for Supplementary Movie 2, except that instead of velocity, the curvature of each point on the simulated contour is now read out relative to its immediate neighbours, as detailed in Methods.

**Supplementary Movie 4. Differential dynamics of nascent adhesions in cells displaying distinct protrusion phenotypes.** B16-F1 control and Lpd KO cells were transfected with EGFP-paxillin, and subjected to time-lapse microscopy during migration on laminin using fluorescence and phase contrast channels, as indicated. A representative B16-F1 control (left panel) is displayed as example for the smooth protrusion phenotype, whereas the middle and right panels show representative examples for nascent adhesion dynamics typically found in protruding cells exhibiting the intermediate and chaotic phenotype, respectively. Phase contrast images (bottom) allow the correlation of distinct protrusion phenotypes with differential adhesion dynamics. The smooth protrusion phenotype is commonly associated with generation of multiple, nascent adhesions, continuously developing distally from previous sets of adhesions during continuous protrusion. Subsets of each population of nascent adhesions then are continuously elongated and developed into mature adhesions. In the intermediate phenotype (middle panels), nascent adhesions are less continuously formed and concentrated at the rear edge of the lamellipodium, frequently coincident with sites of active membrane ruffling. The chaotic protrusion phenotype with its commonly collapsing lamellipodia (right panels) is characterised instead by strongly reduced frequency of nascent adhesion formation.

**Supplementary Movie 5. Dynamics of EGFP-tagged, full length lamellipodin (EGFP-Lpd) in a migrating B16-F1 cell.** B16-F1 control melanoma cell expressing EGFP-Lpd and subjected to fluorescence and phase contrast time-lapse microscopy during its migration on laminin. Note the restriction of EGFP-Lpd localization to the edges of protruding and rearward ruffling lamellipodia, but the lack of association with nascent or mature focal adhesions.

